# OPA1 protects intervertebral disc and knee joint health in aged mice by maintaining the structure and metabolic functions of mitochondria

**DOI:** 10.1101/2024.01.17.576115

**Authors:** Vedavathi Madhu, Miriam Hernandaz-Meadows, Ashley Coleman, Kimheak Sao, Kameron Inguito, Owen Haslam, Paige K Boneski, Hiromi Sesaki, John A Collins, Makarand V. Risbud

**Author notes:** Address correspondence to: Makarand V. Risbud, Ph.D., Department of Orthopaedic Surgery, Thomas Jefferson University, 1025 Walnut Street, Suite 501 College Bldg., Philadelphia, PA 19107, V.M M.V.R. Authors’ contributions: Conceptualization: VM, MVR; Methodology: VM, MVR; Investigation: VM, MHM, AC, KS, KI, OH, PKB, JC; Visualization: VM; Supervision: MVR; Writing - original draft: VM, MVR; Writing - review & editing: VM, KS, MVR, JC, HS.

## Abstract

Due to their glycolytic nature and limited vascularity, nucleus pulposus (NP) cells of the intervertebral disc and articular chondrocytes were long thought to have minimal reliance on mitochondrial function. Recent studies have challenged this long-held view and highlighted the increasingly important role of mitochondria in the physiology of these tissues. We investigated the role of mitochondrial fusion protein OPA1 in maintaining the spine and knee joint health in aging mice. OPA1 knockdown in NP cells altered mitochondrial size and cristae shape and increased the oxygen consumption rate without affecting ATP synthesis. OPA1 governed the morphology of multiple organelles, and its loss resulted in the dysregulation of NP cell autophagy. Metabolic profiling and ^13^C-flux analyses revealed TCA cycle anaplerosis and altered metabolism in OPA1-deficient NP cells. Noteworthy, *Opa1^AcanCreERT2^* mice showed age-dependent disc, and cartilage degeneration and vertebral osteopenia. Our findings suggest that OPA1 regulation of mitochondrial dynamics and multi-organelle interactions is critical in preserving metabolic homeostasis of disc and cartilage.

**Teaser:** OPA1 is necessary for the maintenance of intervertebral disc and knee joint health in aging mice

## INTRODUCTION

Chronic low back pain, often associated with intervertebral disc (IVD) degeneration, and knee and hip joint pain, a common sequela of osteoarthritis, are the leading causes of disability in the aging population worldwide (*1, 2*). In a healthy state, these tissues provide the joint with flexibility and the ability to absorb applied mechanical forces. However, with the onset of degenerative disease, these properties are lost, and the characteristic balance between cell survival, autophagy, and apoptosis becomes dysregulated leading to altered extracellular matrix (ECM) production, and increased tissue catabolism (*2, 3*). Noteworthy, nucleus pulposus (NP) cells of the IVD and chondrocytes of the articular cartilage reside in an avascular, nutrient-limited environment that is hypoxic, aneural, and hyperosmotic. These cells exhibit limited replication and regenerative capacity and robustly express the transcription factor HIF-1α (*4, 5*), and since they possess fewer mitochondria it is thought that they heavily rely on glycolysis to meet their energetic and biosynthetic demands (*6–8*). In contrast to this long-held notion, we recently showed that functional mitochondrial networks exist in NP cells (7), and mounting evidence suggests that mitochondrial dysfunction promotes osteoarthritis development (*2*). The deletion of mitochondrial superoxide dismutase 2 exacerbated, whereas overexpression of catalase or peroxiredoxin 3 reduced the severity of age-associated osteoarthritis in mice, implying that mitochondrial ROS generation plays a role in the pathogenesis of osteoarthritis (*9–11*). Moreover, we have recently shown that to meet metabolic requirements, NP cells evidence an active mitophagic flux governed by the HIF-1α-BNIP3 axis (*6*). In other studies, we demonstrated that the loss of BNIP3 in NP cells resulted in mitochondrial dysfunction, affecting cellular bioenergetics, triggering meta-inflammation, and causing early onset of IVD degeneration (*7*).

As highly dynamic organelles, mitochondria are constantly undergoing fission and fusion that control their mass and numbers. This equilibrium is primarily controlled by the fission protein DRP1 and the fusion proteins MFN1, MFN2, and OPA1 (*12, 13*). The fusion of two adjacent mitochondria promotes membrane tubulation and elongation and is a two-step process: outer mitochondrial membrane (OMM) fusion mediated by mitofusion proteins MFN1 and MFN2 followed by fusion of the inner membrane (IMM) mediated by OPA1 (*14*). During the fusion process, mitochondria control stress and energy needs by incorporating components of the damaged organelles (including mtDNA and mitochondrial respiration complexes) to modulate membrane potential, apoptosis, and calcium signaling (*15*). The cristae of the IMM are the site of the respiratory chain complexes concerned with ATP synthesis and electron transport (*16, 17*).

OPA1 regulates cristae remodeling and apoptosis, independent of mitochondria fusion (*18*). Its alteration or ablation results in cristae disorganization and release of apoptotic cytochrome c (*19*). Conversely, OPA1 overexpression facilitates the formation and stability of respiratory chain supercomplexes (*20*). OPA1 also functions as a sensor of metabolic changes through interactions with IMM solute transporters, facilitating the adaptation of cristae shape and respiration to cellular energy needs (*21*).

Herein we investigated the role of IMM fusion protein OPA1 on IVD and cartilage health. Our findings demonstrate that loss of OPA1 in NP cells results in altered mitochondrial and cristae morphology and mass, with compromised autophagic pathway activity, and metabolism. For the first time, we report the importance of OPA1 in maintaining the morphology of several organelles in the NP cells and demonstrate that conditional deletion of OPA1 (*Opa1^AcanCreERT2^*) causes disc degeneration and osteoarthritis in aged mice. Overall, these findings establish that the IMM fusion protein OPA1 is a critical factor required for maintaining NP and chondrocyte function and promoting the health and function of the IVD and knee articular cartilage.

## RESULTS

### OPA1 is necessary for the preservation of mitochondrial and multi-organelle morphology in NP cells

We investigated the contribution of OPA1 in maintaining mitochondrial morphology and mass by knocking down *Opa1* in primary NP cells using lentivirally delivered ShRNAs. We confirmed OPA1 knockdown by confocal imaging and Western blot analysis (Fig. 1A, A’).

**Figure 1.**
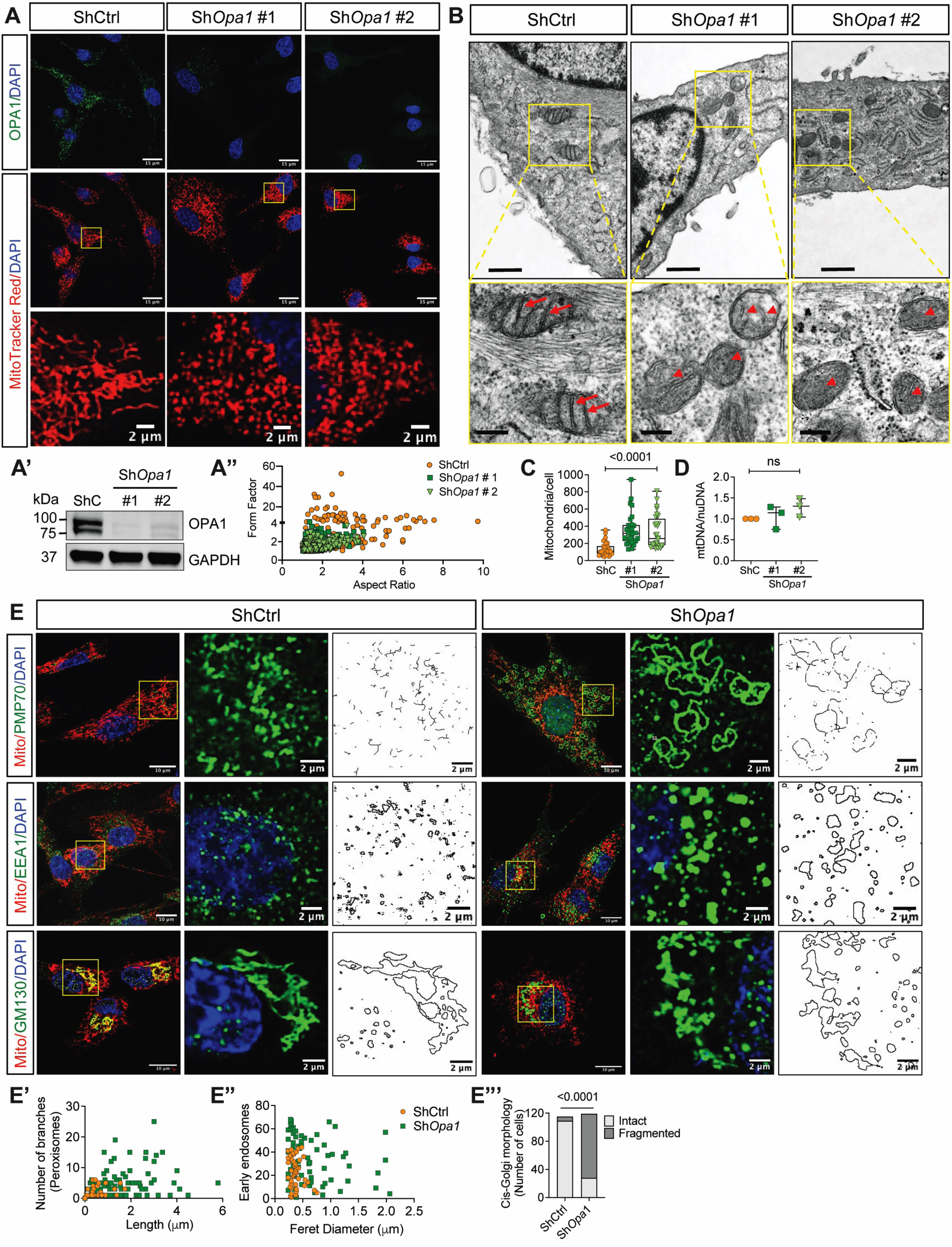
OPA1 is necessary for the preservation of mitochondrial and multiple organelle morphology in NP cells. (A) Immunofluorescence staining for OPA1 and MitoTracker Red in primary NP cells transduced with lentivirally delivered control (ShCtrl) and *Opa1* (Sh*Opa1* #1 and Sh*Opa1* #2) ShRNAs. Scale bar: Top rows- 15 μm, bottom row- 2 μm. (A’) Western blot of OPA1 in NP cells transduced with Sh*Opa1* #1 and Sh*Opa1* #2. (A”) Mitochondrial morphology and network analysis in ShCtrl and Sh*Opa1* NP cells. (B) TEM images of ShCtrl and Sh*Opa1* transduced cells. Scale bar: Top row - 600 nm, bottom row - 200 nm (C) Mitochondrial number measurements from OPA1-deficient NP cells; 30 cells were measured from two independent experiments. (D) The mtDNA content in control and OPA1-deficient cells. Data represented as box and whisker plots showing all data points with median and interquartile range and maximum and minimum values. (E) Immunofluorescence staining for peroxisome marker PMP70, early endosome marker EEA1 and cis-golgi marker GM130. Scale bar: 15 and 2 μm (E’, E”) Morphology analysis of PMP70, EEA1 shows altered morphology of peroxisomes and early endosomes in Sh*Opa1* cells. (E”’) Quantification shows increased fragmented cis-golgi in Sh*Opa1* transduced cells. 100-110 cells were measured for each of the organelle markers. Distribution was performed using chi-square test.

OPA1-deficient NP cells stained with MitoTracker red to visualize mitochondrial morphology showed increased fragmentation, evident from the smaller aspect ratio and form factor of mitochondria (Fig. 1A, A”). TEM imaging showed that mitochondria of the Sh*Opa1* transduced NP cells were smaller in size, with absent or aberrant cristae. Although some cristae were often detached, the tubular structure was preserved with narrow cristae width (Fig. 1B). Notably, OPA1-deficient NP cells also contained more mitochondria than control cells but without changes in mtDNA content, indicating that the increase was due to higher fission and not increased biogenesis (Fig 1C, D). We assessed the effect of OPA1 deficiency on the levels of other fusion and fission proteins. Interestingly, except for the expected decrease in the OPA1, the levels of the outer membrane fusion proteins MFN1, and MFN2 (Fig. S1 A, A’) and the level of the fission protein DRP1 and its receptors FIS1 and MFF were unaffected (Fig. S1 A, A’).

Together these findings showed that OPA1 deficiency affects mitochondrial number, shape, and cristae morphology without compensatory changes in other fusion and/or fission proteins.

We determined the impact of OPA1 deficiency on the morphology of other organelles considering its interaction with many organelles. Using organelle-specific markers, we localized peroxisomes (PMP70), early endosomes (EEA1), late endosomes (RAB7), lysosomes (LAMP1), cis-Golgi (GM130), and trans-Golgi (TGN46). Peroxisomes are vital for detoxification and ß-oxidation of very long-chain fatty acids, which subsequently transports medium-chain fatty acids to mitochondria for further breakdown. Unlike peroxisomes in control cells with lengths between 0.1 to 1 µm, the Sh*Opa1* transduced cells contained hyper-tubulated and hyper-branched peroxisomes that ranged in size from 0.1 to 6 µm (Fig. 1E, E’). We also examined the morphology of early endosomes, which receive cargo and sort it into recycling and degradative compartments. OPA1-deficient NP cells exhibited enlarged endosomes ranging in size from 0.5 - 2 µm, as opposed to endosomes that were smaller than 0.7 µm in control cells (Fig. 1E, E”). We also investigated the morphology of the Golgi complex as the primary secretory pathway organelle. There was a pronounced fragmentation of cis-Golgi in Sh*Opa1* transduced cells (Fig. 1E, E”’, Fig. S2A). In contrast, TGN46-stained trans-Golgi, as well as RAB7-stained late endosomes, and LAMP1-stained lysosomes were unaffected (Fig. S2B). These findings suggest that in addition to maintaining mitochondrial and cristae morphology, OPA1 is essential for sustaining the morphology of organelles that include peroxisomes, early endosomes, and the cis-Golgi in NP cells.

### OPA1 deletion disrupts NP cell autophagy

A striking observation was that OPA1 deficient NP cells showed diminished LC3B-positive autophagosomes (Fig. 2A) while Western blot analysis showed significantly lower LC3II and p62 levels in Sh*Opa1* cells. These findings suggested an overall reduction in autophagy (Fig. 3B, B’). BNIP3 and NIX are BH3-only family proteins that are important in regulating apoptosis, autophagy, and mitophagy. Since we have recently shown that BNIP3 activation and mitochondrial translocation are required for hypoxia-induced mitophagy in NP cells (*6, 22*), we investigated whether BNIP3 mitochondrial localization is affected by OPA1 deletion. Indeed, we observed BNIP3 sequestration in the nucleus of OPA1-deficient cells, without alteration in NIX localization (Fig. S1B, B’) and protein levels (Fig. S1C, C’). The levels of other autophagy-related proteins, BECLIN1, and the ATG12-ATG5 complex remained unaffected whereas those of the apoptotic transcription factor CHOP decreased suggesting Sh*Opa1* cells did not activate apoptosis (Fig. S1C, C’). We examined ubiquitinated protein levels and found no difference between ShCtrl and Sh*Opa1* transduced cells (Fig. S1D, D’). Similarly, levels of PARK2, and its substrate phosphoubiquitin, were unaffected by OPA1 deletion (Fig. 2B, B’, C, C’) (*23*). To ascertain whether OPA1 deletion only impacts autophagy initiation and/or rate of degradation, we treated cells with bafilomycin A1. Cells treated with bafilomycin A1 for 2 hours failed to enhance LC3II accumulation in the Sh*Opa1* cells compared to the ShCtrl, supporting the notion that there was defective autophagic flux (Fig. 2D, D’). The cumulative data indicates dysfunction in the autophagic pathway of OPA1-deficient NP cells.

**Figure 2.**
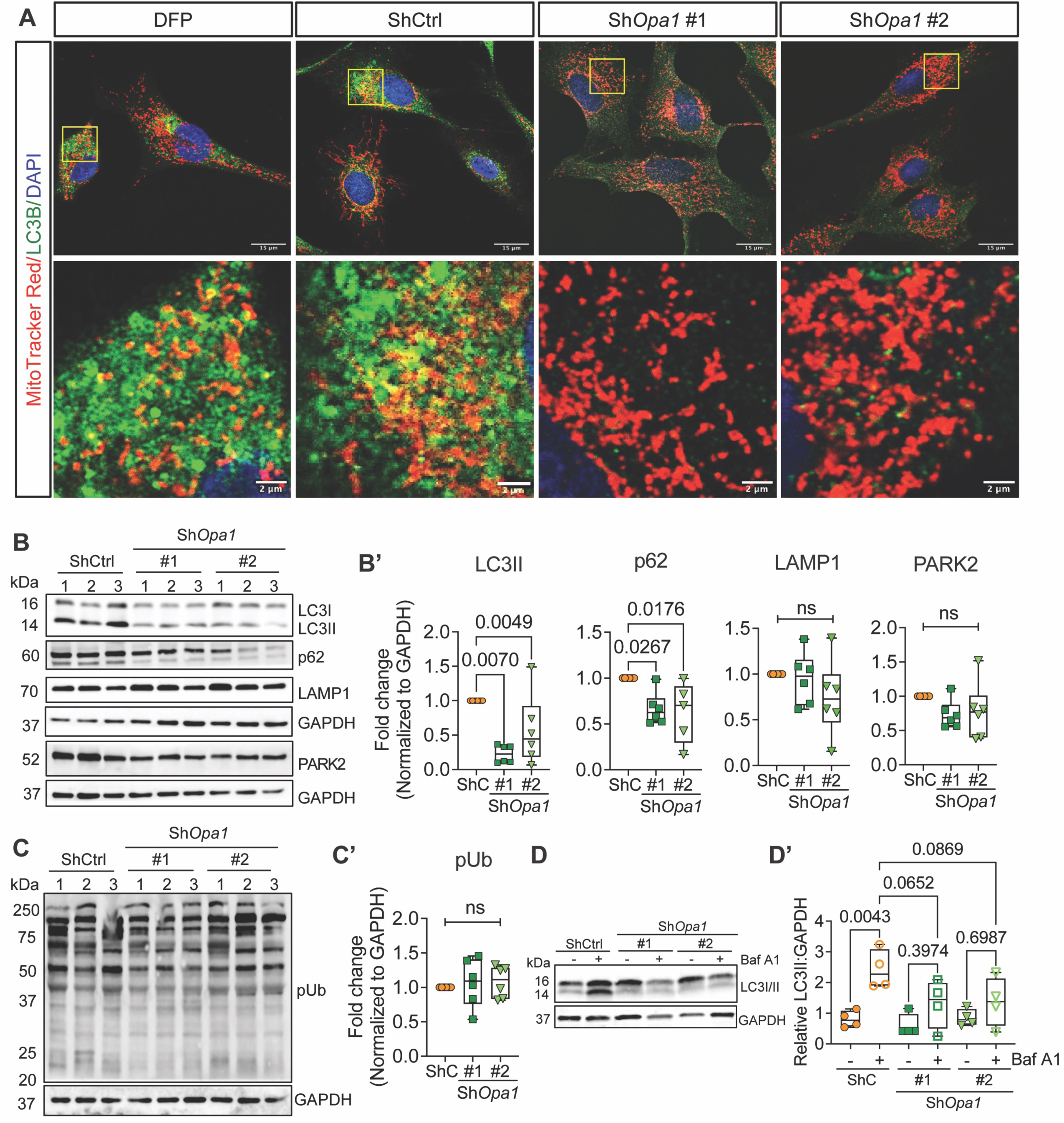
OPA1 deletion disrupts NP cell autophagy. (A) Immunofluorescence staining of LC3B; OPA1-deficient NP cells show a stark absence of LC3B puncta compared to ShCtrl and DFP-treated cells. Scale bar: Top Row-15, Bottom Row - 2 μm. (B, B’) Western blot analysis of 3 independent experiments/group and corresponding densitometric analysis (n = 6 independent experiments) of autophagy/mitophagy pathway markers LC3II, p62, LAMP1, and PARK2 in NP cells sh*Opa1*. (C, C’) Western blot showing 3 representative experiments and densitometry of phospho-ubiquitin after OPA1 deletion (n = 6 experiments). (D, D’) Western blot analysis of LC3B in ShCtrl and Sh*Opa1* transduced cells cultured under hypoxia with or without bafilomycin A (n = 4 experiments). Quantitative data represented as box and whisker plots showing all data points with median and interquartile range and maximum and minimum values. Statistical significance was performed using One-way ANOVA or Kruskal Wallis test with Sidaks’s multiple comparisons test as appropriate.

**Figure 3.**
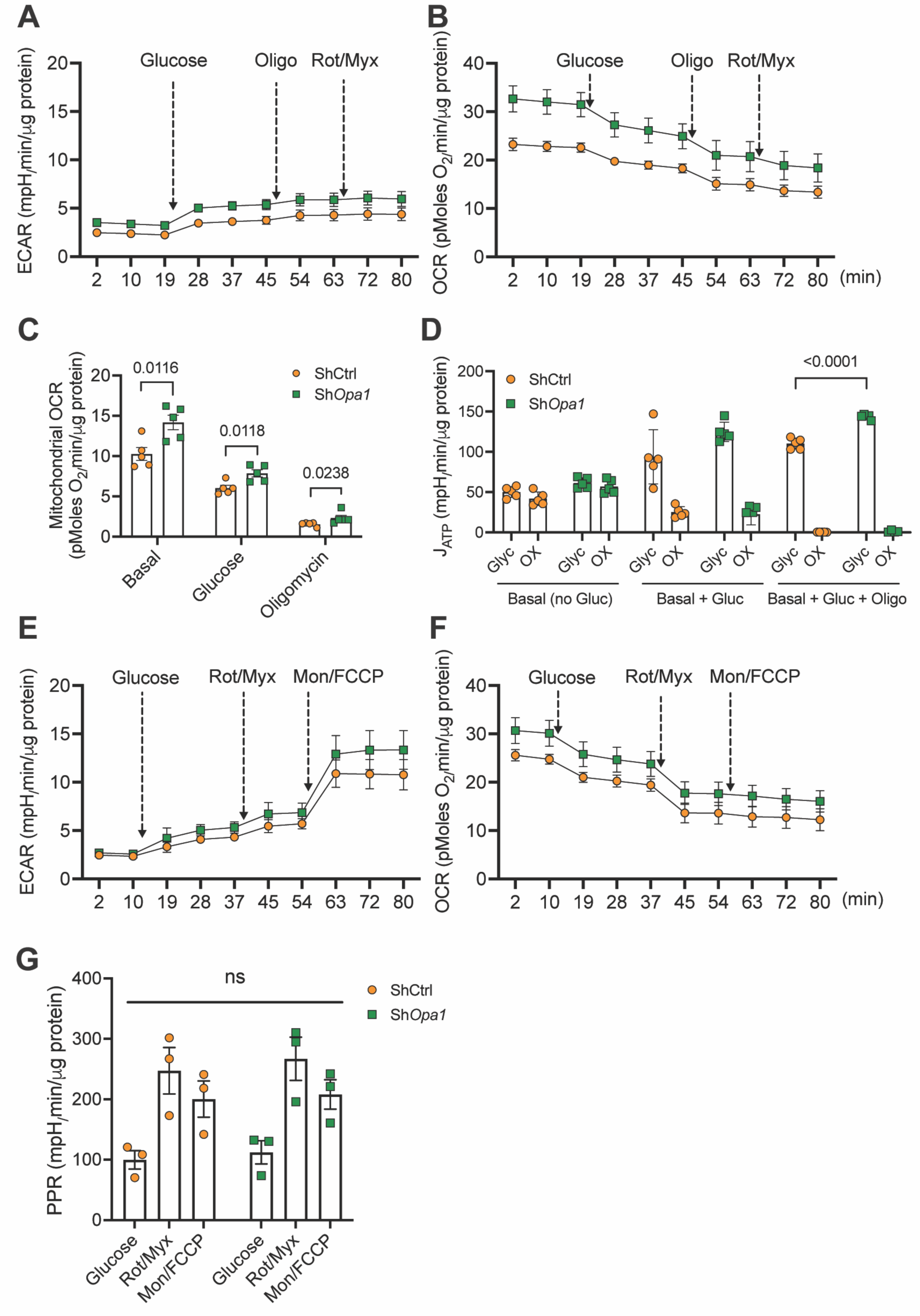
OPA1-deficient NP cells show dysregulated bioenergetics. (A, B) Extracellular acidification rate (ECAR) and oxygen consumption rate (OCR) traces from NP cells transduced with ShCtrl and Sh*Opa1* in the absence of exogenous glucose and after sequential addition of 10 mM glucose followed by oligomycin and rotenone plus myxothiazol. (C) ATP production rate from glycolytic and oxidative metabolism calculated from traces shown in A and B. (D, E) ECAR and OCR traces in the absence of exogenous glucose and after sequential addition of 10 mM glucose followed by rotenone plus myxothiazol and finally monensin plus FCCP. (F) Proton production rate (PPR) of NP cells calculated from traces shown in D and E. Data represent 3-4 independent experiments each with four technical replicates/group. Data is shown mean ± SEM. t-test or Mann-Whitney test or one-way ANOVA was used as appropriate.

### OPA1-deficient NP cells show dysregulated bioenergetics

Since OPA1-deficient NP cells showed mitochondrial fragmentation and abnormal cristae, we assessed their real-time energy metabolism using Seahorse XF Analyzer. The ATP production rates from glycolysis and oxidative metabolism were determined by integrating ECAR and OCR measurements (Fig. 3A, B) (*7, 24*). Under basal conditions, OCR was significantly higher, while ECAR remained unchanged, suggesting OPA1-deficient NP cells consumed more oxygen (Fig. 3 A, B, C). Following the addition of glucose, both ShCtrl and Sh*Opa1* transduced cells showed similar ECAR profiles, but the OCR remained significantly higher and similar to the levels observed under basal conditions (Fig. 3C). Next, we determined levels of ATP generated from glycolysis and oxidative metabolism (Fig. 3D). Glycolytic ATP generation significantly increased in Sh*Opa1* cells; however, oxidative ATP production was constant across groups despite higher oxygen consumption by OPA1-deficient NP cells. These data suggest uncoupling of OXPHOS in OPA1-deficient NP cells, in conjunction with the aberrant cristae morphology. We further investigated the effect of OPA1-deficiency on the glycolytic capacity of NP cells (*7, 25*) by measuring ECAR and OCR profiles under basal (no substrate) conditions and following sequential addition of glucose, rotenone+myxothiazol, and monensin+FCCP (Fig. 3E, F). The Sh*Opa1* cells showed no difference in EACR but demonstrated a prominent increase in OCR traces (Fig. 3E, F). However, the basal glycolytic rate, maximum glycolytic capacity, ATP demand-limited rate, and glycolytic reserve computed from H+/lactate production were unchanged (Fig. 3G). Overall, these results suggest that mitochondrial fragmentation and disorganized cristae in OPA1-deficient cells alters energy metabolism of NP cells.

### OPA1-deficient NP cells evidence altered glucose and glutamine metabolism

To understand the broader metabolic alterations caused by OPA1 deficiency, we performed widely targeted metabolic profiling on NP cells. A total of 261 metabolites that satisfied the QC limit of < 30% coefficient of variation (CV) were imported individually into the SIMCA-p program for multivariate analysis. An unsupervised principal component analysis (PCA) and supervised partial least square-discrimination analysis (PLS-DA) models were established which showed a clear separation between the Sh*Opa1* and ShCtrl samples (Fig. 4A, B). We identified 36 downregulated and 9 upregulated metabolites (FDR ≤ 0.05) in OPA1-deficient cells (Fig. 4C). Concerning altered metabolites, the levels of glycolytic metabolite glucose-6-phosphate and TCA metabolite malate, as well as the collagen-related amino acids hydroxyproline, and hydroxylysine were raised (Fig. 4D-G), whereas fatty acid metabolites stearic acid and acetylcarnitine and the one carbon metabolites s-adenosyl homocysteine and s-adenosyl methionine were significantly lowered in OPA1 knockdown cells. (Fig. 4H-K). Additionally, the Sh*Opa1* group showed a significant increase in NADP with lower AMP, without affecting the overall ADP and ATP levels (Fig. 4L-O). However, AMP/ATP and ADP/ATP ratios were affected in knockdown NP cells (Fig. 4P, 4Q). Analysis of upregulated metabolites revealed enrichment in galactose metabolism, nucleotide sugar metabolism, starch, and sucrose metabolism, ascorbate and aldarate metabolism, pentose and glucuronate interconversion, sphingolipid metabolism, pentose phosphate pathway activity, glycolysis/gluconeogenesis, and inositol phosphate metabolism-related metabolites. Similarly, metabolic pathway analysis of upregulated entities showed nucleotide sugar metabolism, malate-aspartate shuttle activity, transfer of acetyl group into mitochondria, and starch and sucrose metabolism, and lactose synthesis were the most impacted pathways in Sh*Opa1* transduced cells (Fig. 4R, Fig.S3A).

**Figure 4.**
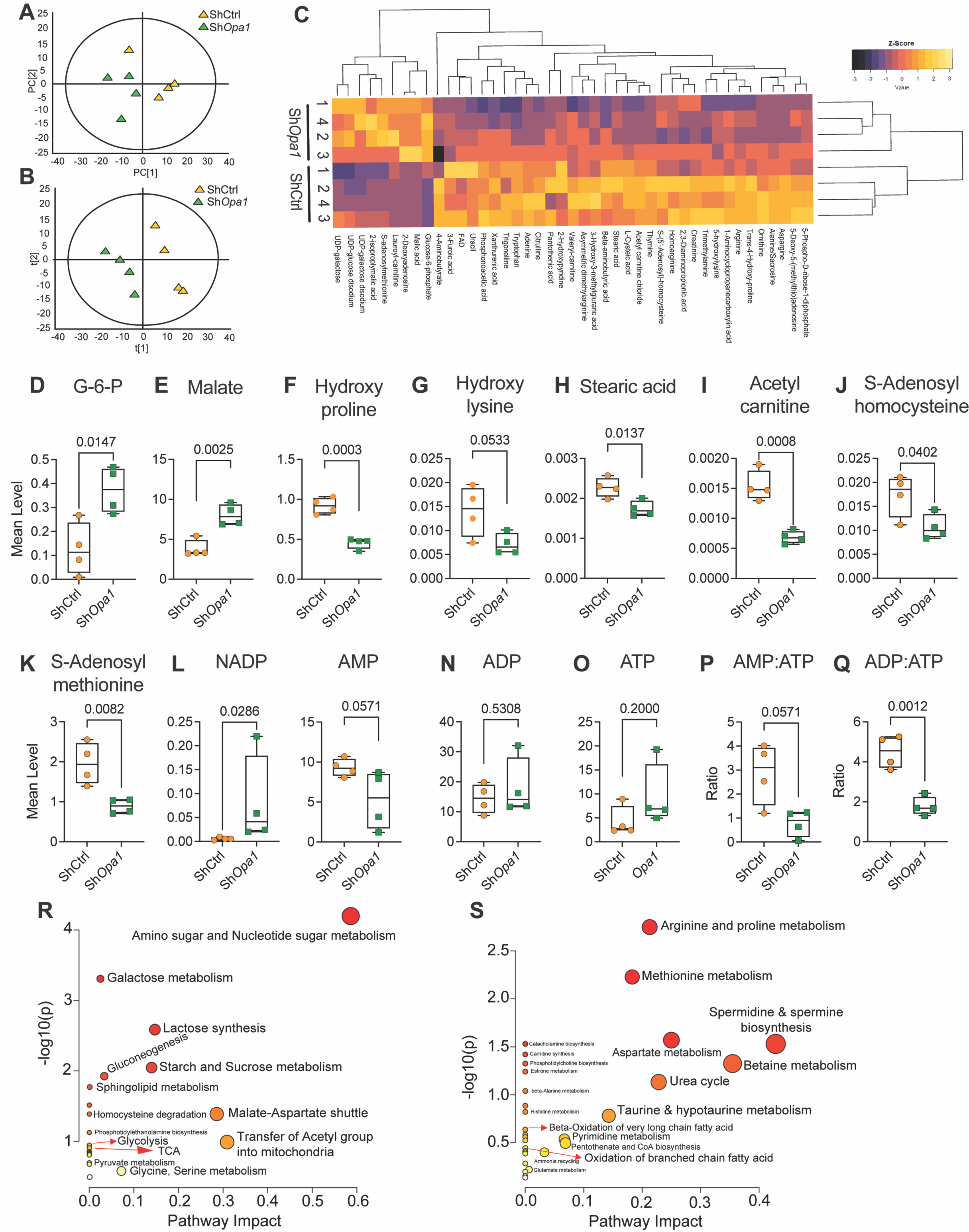
OPA1 is an important regulator of hypoxic NP cell metabolism. (A) Unsupervised principal component analysis (PCA) and (B) Supervised partial least square-discrimination analysis (PLS-DA) model of widely targeted small metabolites from NP cells transduced with ShCtrl and Sh*Opa1*. n = 4 independent experiments (C) Heat map of normalized concentration of metabolites differentially present between ShCtrl and Sh*Opa1* using FDR adjusted p-value ≤0.06%. (D-Q) Mean levels of a select group of measured metabolites from ShCtrl and Sh*Opa1* NP cells (D) Glucose-6-phosphate, (E) Malate, (F) Hydoxyproline, (G) Hydroxylysine, (H) Stearic acid, (I) Acetylcarnitine, (J) S-adenosylhomocysteine and (K) S-adenosylmethionine, (L) NADP, (M) AMP, (N) ADP, (O) ATP, (P) ADP/ATP and (Q) AMP/ATP. (R, S) Metabolite set enrichment analysis (MSEA) showing pathway impact and p-value of significantly upregulated and downregulated metabolites (FDR≤0.05) from Sh*Opa1*transduced NP cells. Quantitative data is represented as box and whisker plots showing all data points with median and interquartile range and maximum and minimum values. Statistical significance was computed using t-test or Mann-Whitney test as appropriate (D-Q).

Arginine, cysteine, methionine, proline, alanine, aspartate, and glutamate were among the enriched metabolites that were significantly downregulated, and the enrichment analysis showed downregulation of aminoacyl-tRNA biosynthesis, pantothenate and CoA biosynthesis, pyrimidine and purine metabolism and taurine and hypotaurine metabolism in Sh*Opa1* cells.

Likewise, the most impacted downregulated pathways in Sh*Opa1* cells were spermidine and spermine biosynthesis, and betaine, urea, aspartate, methionine, arginine, and protein, taurine and hypotaurine, pyrimidine, and pantothenate and CoA metabolism, and ammonia recycling (Fig. 4S, Fig. S3B).

To further delineate the utilization of major metabolic substrates glucose and glutamine, OPA1-silenced NP cells were cultured for 24 hours under hypoxia with a 50% enrichment in either [1,2]-^13^C-glucose or U^13^C-glutamine (Fig 5A). Based on the lactate and glutamate of the medium we calculated the glycolytic, pentose cycle, PDH, PC, PDH+PC, and PHD/PC flux, as previously described (*7*). OPA1-silencing did not affect glucose flux through glycolysis or the pentose cycle (Fig. 5B, C). ^13^C enrichment in medium glutamate provided a measure of glucose flux into the TCA cycle via PDH and PC. PDH flux was significantly decreased, without altering flux through PC; combined PDH+PC flux or the PDH/PC also remained unchanged (Fig. 5D-G). Analysis of extracted metabolites from the cell pellet showed a modest increase in sigma mean (∑ mn = 1 *M1 +2*M2+3*M3, etc.) of alanine (m/z 260) and a decrease in serine (m/z 390) with minimal changes in lactate (m/z 261), citrate (m/z 591), and succinate (m/z 289). A small decrease in M1 glutamate (m/z 432) as well as enhanced enrichment in palmitate (m/z 313) and stearic acid (m/z 341) was also noted (Fig. 5H-O). When U^13^C-glutamine utilization was measured, Sh*Opa1* cells evidenced a reduction in TCA cycle intermediates citrate (m/z 591), succinate (m/z 289), fumarate (m/z 287), malate (m/z 419), and aspartate (m/z 418), without changes in the enrichment of lactate (m/z 261) (Fig. 5P-U). Both hypoxia and ETC inhibition increased the levels of succinate (*26*). Interestingly, however, our ^13^C-MFA data showed increased succinate ∑ mn compared to fumarate in both ShCtrl and Sh*Opa1* cells, suggesting involved impairment of the oxidative TCA cycle. Furthermore, when we assessed succinate oxidation (fumarate M+4/succinate M+4) and fumarate reduction (succinate M+3/fumarate M+3), changes in the oxidative process were apparent, albeit a downward trend in fumarate reduction in Sh*Opa1* cells (Fig. 5V, W). These studies lent considerable support for the notion that OPA1 is required for optimal NP cell metabolism.

**Figure 5.**
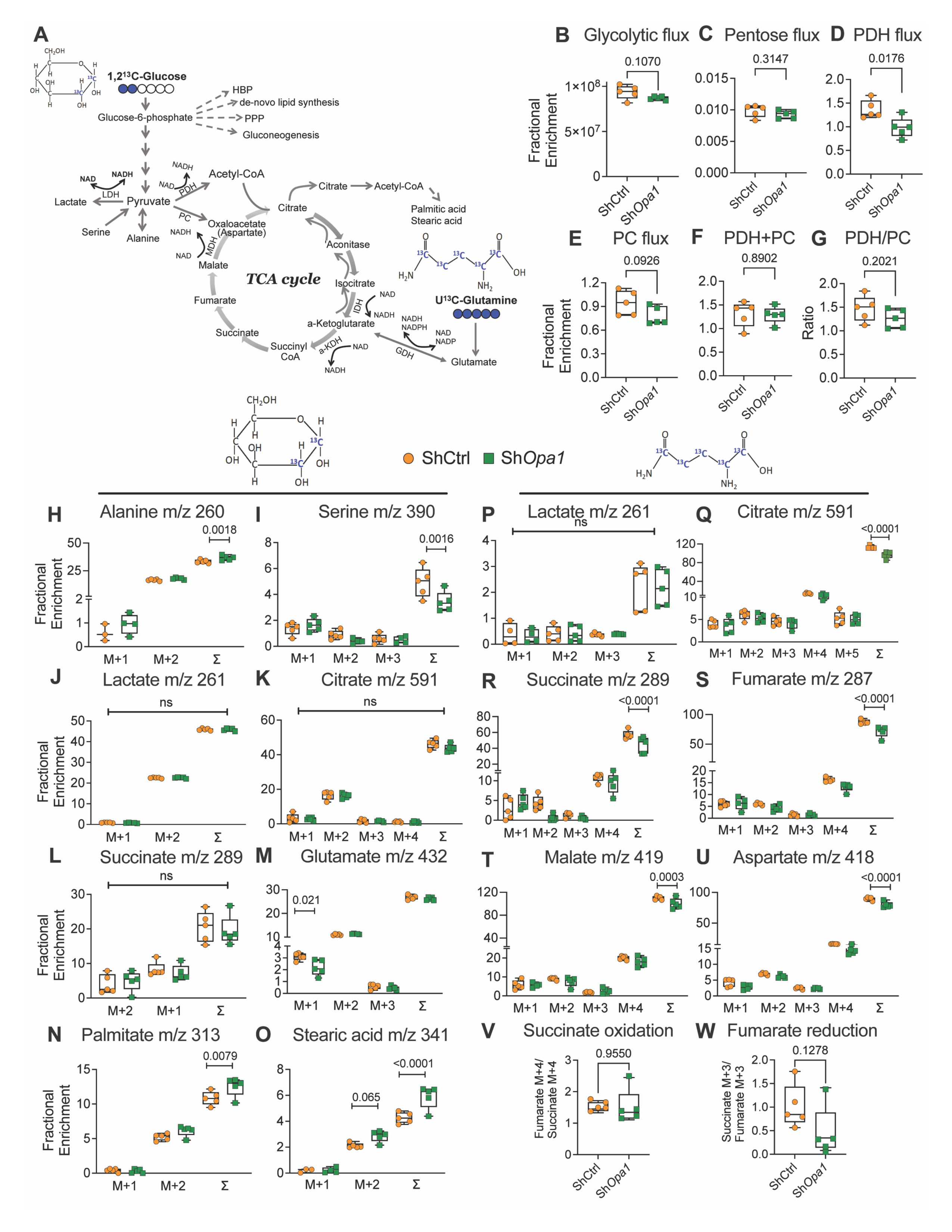
OPA1-deficient NP cells evidence altered glucose and glutamine metabolism. (A) Summation of flux results through glycolysis, pentose, and TCA cycle using [1,2]-^13^C-glucose and U^13^C-glutamine. (B-G) [1,2]-^13^C-glucose enrichment in the culture media from NP cells transduced with ShCtrl and Sh*Opa1* and cultured under hypoxia for 24 h. (B) Glycolysis flux as measured by enrichment of M2 lactate m/z 91, (C) Pentose cycle flux as measured by lactate (M1/M2)/ (3 + M1/M2) (D) PDH flux as measured glutamate m/z 103, (E) PC flux measured as glutamate m/z 104 (F) PC+PDH flux measured as glutamate m/z 131, (G) PDH/PC flux as measured by glutamate m/z 103-M1/104-M2. (H-O) [1,2]-^13^C-glucose tracing measured from the cell pellets into metabolites (H) alanine m/z 260, (I) serine m/z 390, (J) lactate m/z 261, (K) citrate m/z 591, (L) succinate m/z 289, (M) glutamate m/z 432, (N) palmitate (C16:0) m/z 313, (O) stearic acid (C18:0) m/z 341. (P-W) U^13^C-glutamine tracing measured from cell pellets of NP cells transduced with ShCtrl and Sh*Opa1* and cultured under hypoxia for 24 h (P) lactate m/z 261, (Q) citrate m/z 591, (R) succinate m/z 289, (S) fumarate m/z 287, (T) malate m/z 419, (U) aspartate m/z 418, (V) succinate oxidation and (W) fumarate reduction. Quantitative data is represented as box and whisker plots showing all data points with median and interquartile range and maximum and minimum values. Data points with negative values (undetectable ∑ mn) are not included. Statistical significance was computed using t-test or Mann-Whitney test as appropriate.

### Conditional deletion of *Opa1* in IVD accelerates age-associated degeneration

To date, the *in vivo* role of OPA1 in the spine has not been described. To gain a better understanding of how OPA1 impacts spinal health, we generated *Opa1* conditional knockout mice by administering tamoxifen to 3-month-old Acan^CreERT2^*Opa1^fl/fl^* (*Opa1*cKO) and *Opa1^fl/fl^*(WT) mice (Fig. 6A, B) (*27*). In adult mice, the Acan-Cre^ERT2^ driver is highly effective in targeting all three compartments of the IVD as well as the articular and growth plate cartilages (*28*). The successful deletion of OPA1 was confirmed by mRNA and protein evaluation of the NP and annulus fibrosus (AF) tissues (Fig. 6C, D). We performed a quantitative histopathological analysis of IVD morphology using the Modified Thompson score on 7, and 12- month-old *Opa1*cKO mice (Fig. S4) indicated no noticeable degeneration in the IVD in the lumbar (Fig. S4. A-A”, B-B”) or caudal regions of the spine (Fig. S4. C-C”, D-D”) compared to WT mice. At 12 months, however, caudal discs of *Opa1*cKO mice showed changes in NP cell morphology and AF hyperplasia (Fig. S4D”’). When *Opa1*cKO mice were evaluated at 20 months, a prominent degenerative phenotype in the NP and AF tissues of caudal IVDs was evident (Fig. 6E). The distribution of Modified Thompson grading scores showed a significantly higher proportion of *Opa1*cKO discs had NP and AF compartments scores of 3 or 4, indicating severe degeneration (Fig. 6E’, E”). At 20 months, the degenerative phenotype was characterized by a diminished SafraninO stained NP extracellular matrix, a significant loss of NP cells with the remainder of cells acquiring a hypertrophic chondrocyte-like morphology and irregularities in AF lamellar organization (Fig. 6E). There was also a clear loss of demarcation between NP and AF tissue boundaries in *Opa1*cKO mice (Fig. 6E). Those discs that retained notochordal NP cell bands displayed morphological changes such as loss of cytosolic vacuoles (Fig. 6E”’). In addition to these morphological changes, *Opa1cKO* mice exhibited AF hyperplasia, and this was reflected in increased tissue area (Fig. 6F, F’). Picrosirius red staining coupled with polarized light imaging was used to ascertain if there were alterations in collagen matrix organization (Fig. 6G). At 20 months, unlike WT mice, *Opa1*cKO mice showed the presence of collagen fibers in the NP compartment. Moreover, when the fraction of fibrous tissue area in the NP was measured; *Opa1*cKO animals also exhibited a higher proportion of fibrous tissue area than a few discs in WT mice that showed NP fibrosis (Fig. 6G’). In the AF tissue a higher proportion of thin collagen fibers than WT mice suggesting increased collagen turnover in the *Opa1*cKO mice (Fig. 6G”, G”’). Notably, compared to caudal discs, lumbar IVDs showed a milder phenotype; there was a higher proportion of discs with NP and AF compartments scoring grade 4 but when scores of all discs were averaged it did not reach statistical significance (Fig. S5A, A’, A”). Similar to caudal IVDs, the NP compartment of lumbar IVDs in 20-month-old *Opa1*cKO mice displayed a higher prevalence of fibrous tissue compared to WT mice indicating degenerative changes (Fig. S5B, B’).

**Figure 6.**
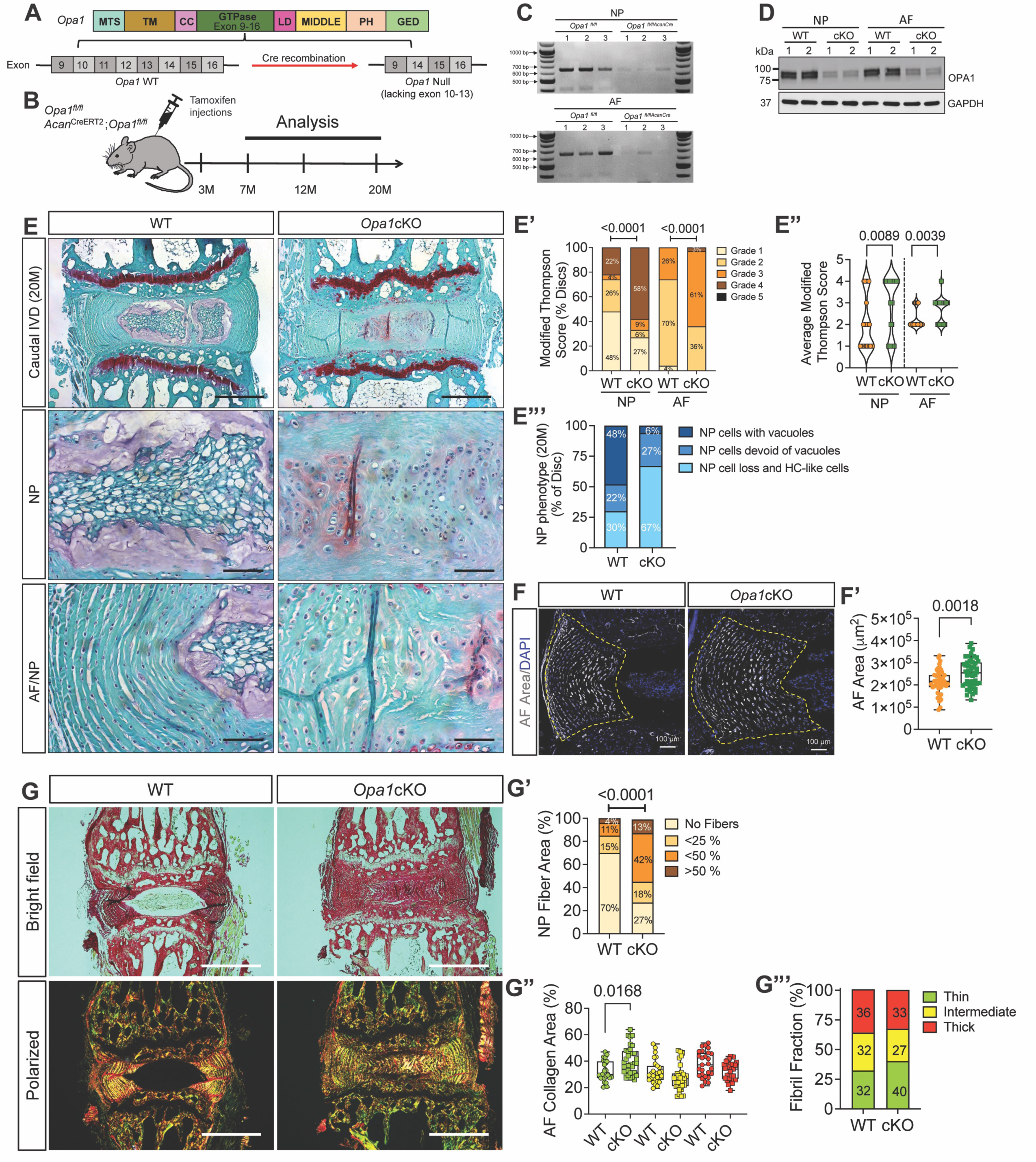
Conditional deletion of *Opa1* in IVD accelerates age-associated degeneration. (A) Schematic showing tamoxifen induced *Acan^CreERT2^*mediated deletion of exons 10-13 of *Opa1* to generate *Opa1* null allele. (B) Tamoxifen treatment and analysis timeline of WT (*Opa1*^fl/fl^) and *Opa1^AcanERT2^* (*Opa1*cKO) mice. (C) RT-PCR analysis shows *Opa1* deletion in NP and AF tissues of *Opa1*cKO mice (n=3 mice/genotype). (D) Western blotting of NP and AF tissue protein from *Opa1*cKO mice shows a significant decrease in OPA1 levels (n=2 mice/genotype). (E) Representative Safranin O/Fast Green/Hematoxylin staining of 20-month-old WT and *Opa1*cKO caudal disc sections showing overall morphology of disc, NP, AF compartments, and NP/AF interface. Scale bar: Top row-500 μm, Bottom rows - 100 μm. (E’, E”) Modified Thompson Scores of NP and AF compartment of WT and *Opa1*cKO caudal discs. (E”’) Distribution graph showing NP phenotype of 20-month-old WT and *Opa1*cKO. n = 9 WT (4F, 5M), 11 *Opa1*cKO (6F, 5M) mice; 3 caudal discs/animal, 27-33 discs/genotype. (F, F’) Dotted lines demarcate AF compartment hyperplasia in *Opa1*cKO caudal discs and compartment area. scale bar = 100 μm. (G) Representative Picrosirius red stained brightfield and polarized light images of 20-month-old mice caudal discs show evidence of NP collagen fibers. (G’) The distribution of discs based on %NP are occupied by collagen fibers. (G”, G”’) Quantitative analysis of thin, intermediate, and thick AF collagen fibril fraction. n = 9-11 mice/genotype; 3 caudal discs/animal, 27-33 discs/genotype. Significance for E’, E”, G’, G”’ was determined using a chi-square test. The significance of the fiber percentage of AF collagen area (G”) was determined using Kruskal- Wallis with Dunn’s test. Quantitative Data represents violin (E’) or box and whiskers (F’, G”) plot showing all data points with median and interquartile range and maximum and minimum values. Significance was determined using an unpaired t-test or Mann-Whitney test, as appropriate.

### The lack of OPA1 affects NP cell phenotype and alters the IVD matrix composition

We assessed the NP cell phenotype by measuring the abundance of the phenotypic markers, carbonic anhydrase (CA3), and glucose transporter I (GLUT1) in 20-month-old mice. Strikingly, expression of these markers was lost by the few cells that remained in the NP compartment, suggesting the native notochord cell population underwent a phenotypic switch (Fig. 7A, B).

**Figure 7.**
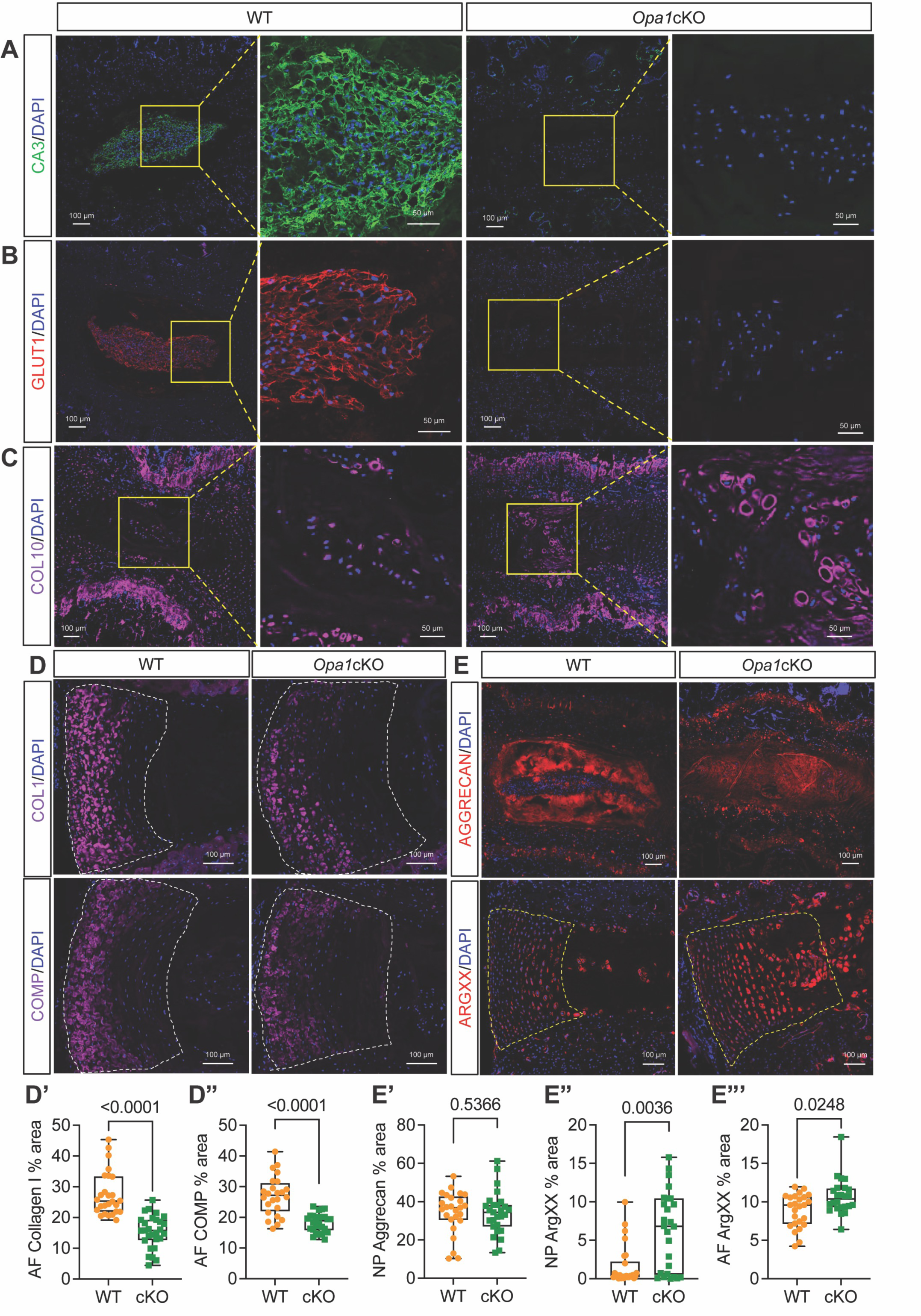
The OPA1-deficiency affects NP cell phenotype and alters the IVD matrix composition. Qualitative immunohistological staining of 20-month WT and *Opa1*cKO caudal discs for (A) carbonic anhydrase 3 (CA3); (B) glucose transporter 1 (GLUT1) and (C) collagen 10 (COLX). Scale bar = 100 and 50 μm. Qualitative and quantitative immunohistological staining for (D, D’) collagen 1 (COL1); (D, D”) cartilage oligomeric matrix protein (COMP), (E, E’) aggrecan (ACAN); (E, E”) aggrecan G1 neoepitope (ARGxx), a marker of aggrecan degradation. Scale bar = 100 μm. (n = 9-11 mice/genotype, 1-3 discs/mouse, 13-14 discs/genotype/marker). Quantitative measurements are shown as Box and whisker plots showing all data points with median and interquartile range and maximum and minimum values. Significance was determined using an unpaired t-test or Mann-Whitney test, as appropriate.

In the small number of IVDs that retained the NP cell band, the expression of these phenotypic markers was evident. Considering the NP cells may have undergone a phenotypic shift, we stained IVD sections for COLX, a marker of hypertrophic chondrocytes. Staining was considerably higher in both the NP and AF of *Opa1*cKO mice (Fig. 7C), suggesting that resident cells in the NP compartment acquired hypertrophic chondrocyte-like characters. The ECM composition of the *Opa1cKO* was also determined. *Opa1*cKO had a lower AF abundance of collagen I (COLI) (Fig. 7D, D’) and cartilage-oligomeric matrix protein (COMP), an important non-collagenous matrix component (Fig. 7D, D”). While ACAN staining revealed no major differences across genotypes (Fig. 7E, E’), the NP and AF compartments of *Opa1*cKO mice displayed a large increase in ARGxx, an ACAN neoepitope formed by ADAMTS4/5 dependent cleavage (Fig. 7E, E”). Overall, these findings suggest that OPA1 is required for maintaining the IVD cell phenotype and major structural components of the ECM.

### *Opa1*cKO mice evidence alterations in vertebral bone health and disc height index

To investigate the impact of OPA1 deletion on vertebral bone health of the aging spine we performed micro-computed tomography (μCT) on the lumbar (L3-6) vertebrae of mice. In WT mice, three-dimensional (3D) reconstructions showed expected trabecular bone loss with aging; however, *Opa1*cKO mice exhibited a significant reduction in vertebral trabecular bone architecture at 7- and 12-months (Fig. 8A). Thus, bone volume/trabecular volume (BV/TV), bone mineral density (BMD), trabecular thickness (Tb.Th), and trabecular number (Tb.N) was decreased. The change in bone structure in the 7 and 12-month-old *Opa1*cKO was maintained at 20 months implying early onset osteopenia (Fig. 8B-E). As expected, trabecular separation was also greater in 7-month-old *Opa1*cKO mice (Fig. 8F); however, at 7 and 12 months, the structural model index (SMI), a parameter that identifies the rod-like structure of trabeculae and is linked to bone strength and fracture risk, was considerably greater (Fig. 8G). Interestingly, unlike trabecular bone, changes in cortical bone structure were evident in 20-month-old *Opa1*cKO mice, with a substantial increase in mean total cross-sectional bone area (B.Ar.), tissue area (T.Ar.) and the cross-sectional thickness (Cs.Th.), without a change in tissue mineral density (TMD) (Fig. 8H-L). We also noted the increased vertebral length and disc height index (DHI) in *Opa1*cKO mice at 7- and 12- months (Fig. 8M-P), correlated with IVD degeneration. These findings suggested that the loss of OPA1 in *Acan*-expressing growth plate chondrocytes altered bone structural parameters, implying that the mitochondrial fusion protein OPA1 plays an important role in the control of vertebral bone health.

**Figure 8.**
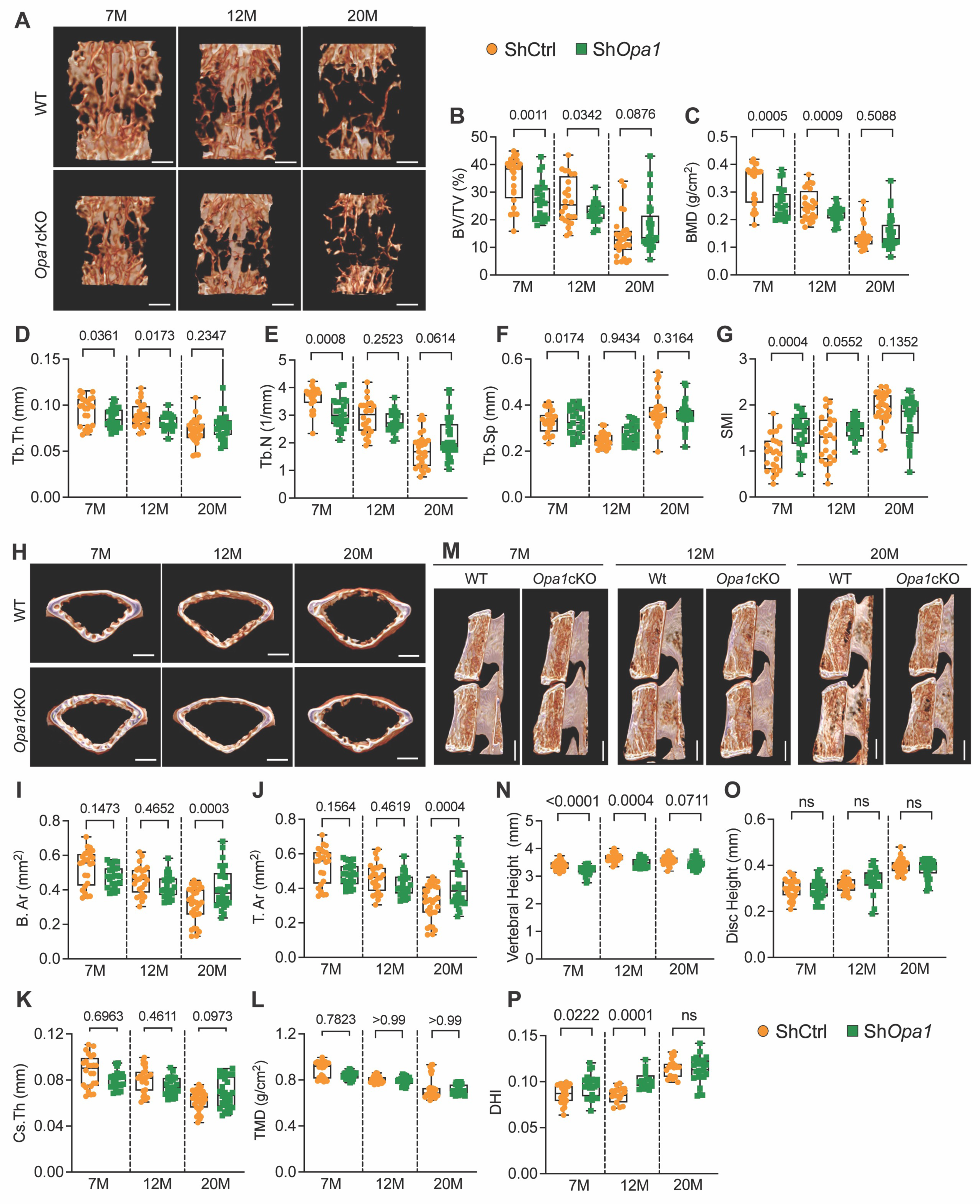
*Opa1*cKO mice evidence alterations in vertebral bone health and disc height index. (A) The representative 3D reconstruction of hemi-section and trabecular lumbar vertebral bone from 7-month, 12-month, and 20-month-old WT and *Opa1*cKO mice. Scale bars 500 μm. Vertebral trabecular bone parameters (B) Percent bone volume/ tissue volume (BV/TV) (%), (C) Bone mineral density (BMD) (g/cm^3^), (D) trabecular thickness (Tb.Th) (mm), (E) trabecular number (Tb.N) (1/mm), (F) trabecular separation (Tb.Sp) (mm), (G) structural model index (SMI) of WT and *Opa1*cKO mice. (H) Representative 3D reconstruction of transverse section through vertebrae of WT and *Opa1*cKO mice showing cortical shell geometry and corresponding quantitative parameters (I) mean total cross-sectional thickness bone area (B.Ar) (mm^2^), (J) mean total cross-sectional tissue area (T.Ar) (mm^2^), (K) cross-sectional thickness (Cs.Th) (mm), (L) Tissue mineral density (TMD) (g/cm3). Scale bars 500 μm. (M) Representative microCT reconstructions of hemi-sections through vertebrae of WT and *Opa1*cKO mice to determine (N) Vertebral length (VL) (mm), (O) disc height (DH) (mm), and (P) disc height index (DHI). Scale bars 1 mm. Quantitative data shown as box and whisker plots showing all data points with median and interquartile range and maximum and minimum values. 7-month n = 10 WT (3F, 7M), 8 *Opa1*cKO (3F, 5M); 12-month n = 10 WT (6F, 4M), 10 *Opa1*cKO (4F, 6M); 20-month n = 9 WT (4F, 5M), 11 *Opa1*cKO (6F, 5M), 4 vertebrae and 3 discs/mouse were analyzed. Significance was determined using t-test or Mann-Whitney test, as appropriate.

### *Opa1*cKO mice show increased severity of age-associated OA

Since the *Acan^CreERT2^*allele also efficiently targets articular cartilage, we investigated if OPA1 deletion has deleterious effects on knee articular cartilage and overall knee joint health in mice.

μCT images showed evidence of bone spurs in the knee joints of 20-month but not 12-month-old Opa1cKO mice (Fig. 9A, Fig. S6A). μCT was also used to evaluate tibial subchondral bone volume fraction (BV/TV), trabecular thickness (Tb.Th), trabecular separation (Tb.Sp), and subchondral bone plate thickness (SCBP) in the medial and lateral tibial plateaus. These analyses showed a significant increase in subchondral bone thickness in the lateral compartment of the 20- month-old mice (Fig. 9B-E); none of the measured bone metrics showed changes at 12 months (Fig. S6B-E). To study cartilage structure, we stained 12- and 20-month-old knee joints sectioned in the mid-coronal plane with H&E and toluidine blue as we have previously described (*29*) (Fig. 9F, G). At 20 months, *Opa1*cKO mice showed severe OA with loss of articular cartilage on the lateral tibial plateaus and femoral condyle, but only minor degradation of articular cartilage in the medial compartment (Fig. 9F). Furthermore, H&E staining revealed osteophyte formation in the medial compartment of WT mice, and osteophyte formation in both the lateral and medial compartments of *Opa1*cKO animals (Fig. 9G). When these structural changes were quantified, *Opa1*cKO mice exhibited a significant increase in articular cartilage score (ACS) in the lateral compartment of the knee at 20-months. However, there were no deviations in toluidine blue scores and osteophyte scores between compartments or genotypes (Fig. 9H-J). However, the cumulative osteophyte score in *Opa1*cKO mice was significantly higher than in WT (Fig. 9K).

**Figure 9.**
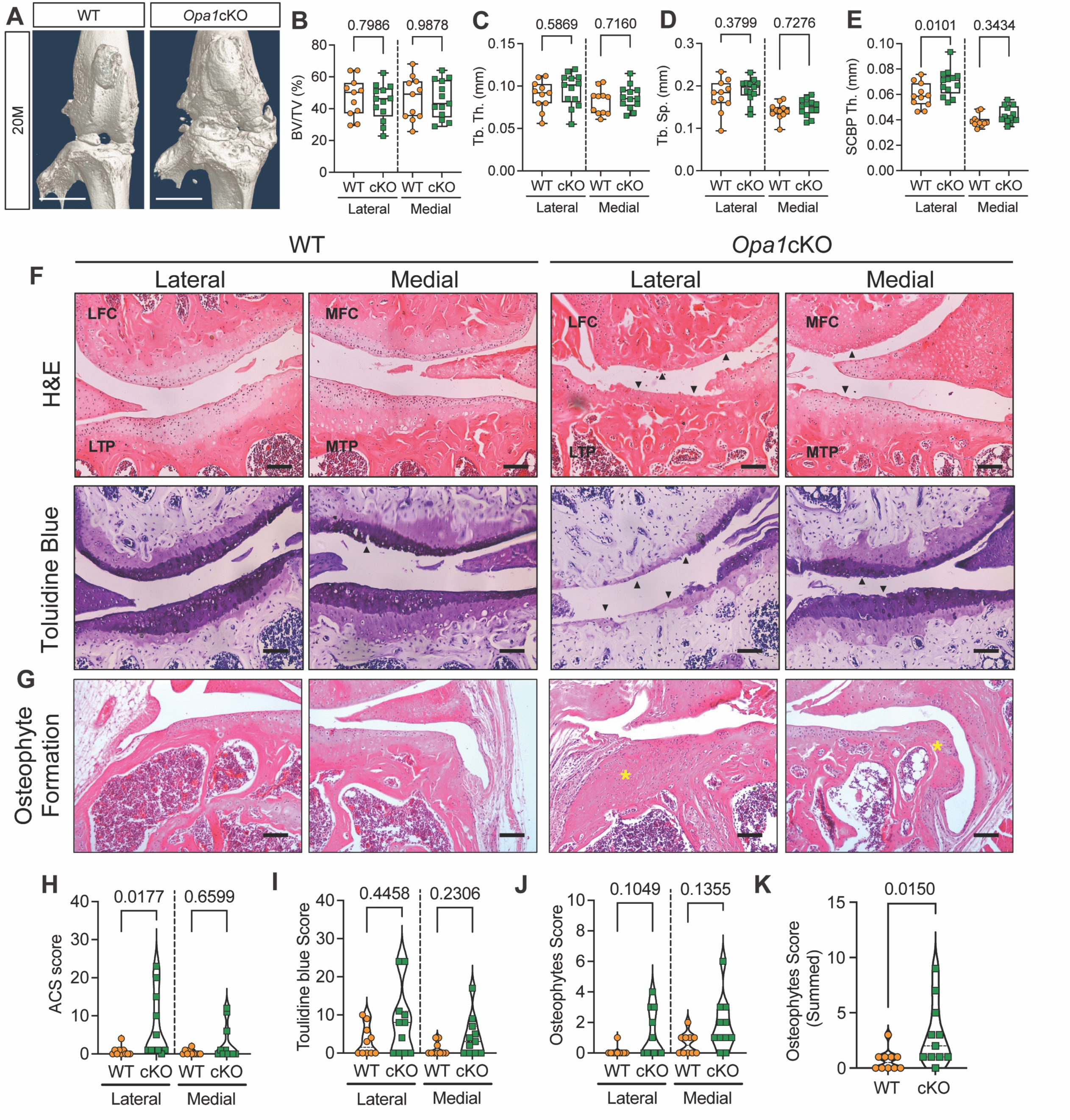
*Opa1*cKO mice show increased severity of age-dependent OA. (A) Representative 3D reconstruction of the whole knee joint of 20-month-old WT and *Opa1*cKO mice. Extensive bone spurs are evident in the *Opa1*cKO joint. Scale bar: 500 μm. (B-E) Quantitative analyses of tibial subchondral bone parameters on the medial and lateral tibial plateaus of 20-month-old WT and *Opa1*cKO mice (B) bone volume fraction (BV/TV), (C) trabecular thickness (Tb.Th), (D) trabecular separation (Tb.Sp), (E) subchondral bone plate thickness (SCBP.Th). n = 11 WT (4F, 6M); 13 *Opa1*cKO (5F, 8M). (F) Representative images of H&E and toluidine blue stained midcoronal sections showing the lateral and medial tibial plateaus and femoral condyles from WT and *Opa1*cKO mice. Extensive cartilage damage (black arrowheads) predominantly in the lateral and medial compartments of *Opa1*cKO mice. Scale bar: 100 μm. (G) Representative images of H&E stained sections showing the presence of large osteophytes on the medial and lateral tibial plateaus of *Opa1*cKO mice (yellow asterisks). Scale bar 100 μm. (H-J) Summed medial (MTP, MFC) and lateral (LTP, LFC) scores for (H) Articular Cartilage structure (ACS), (I) Toluidine blue (J-K) osteophytes. LFC = lateral femoral condyle, LTP = lateral tibial plateau, MFC = medial femoral condyle, MTP = medial tibial plateau. n = 10 WT (4F, 6M), n = 11 *Opa1*cKO (5F, 6M). Data is represented as box and whiskers (B-E) and violin (H-K) plots showing all data points with median and interquartile range and maximum and minimum values. Significance was determined using t-test or Mann-Whitney test as appropriate.

Moreover, H&E staining of joints from 20-month-old *Opa1*cKO mice also showed synovial hyperplasia/ossification in the lateral and medial compartments (Fig. 10A) however, there were no morphological changes between WT and *Opa1*cKO mice at 12 months (Fig. S6. F, G). Additionally, we conducted histomorphometric analysis on both lateral and medial knee joint compartments of 12 and 20-month-old mice. In the lateral compartment of 20-month *Opa1*cKO mice there was a significant decrease in the area and thickness of articular cartilage and calcified cartilage (Fig. 10B, B’, C, C’, D, D’). SCBP area and thickness were significantly increased in *Opa1*cKO mice when compared to controls, indicating enhanced subchondral bone sclerosis in OPA1 loss mice, a finding that aligns with our µCT data (Fig. 10D, D’). We also noted a considerable increase in the synovial hyperplasia/ossification score in the lateral joint compartment of 20-month-old *Opa1*cKO mice (Fig. 10E). On the other hand, 12-month-old mice showed no significant morphological and histomorphometric changes or expression of an OA phenotype (Fig. S6F-M’). It is interesting to note that the profound OA phenotype in *Opa1*cKO mice was primarily observed in the lateral compartment as opposed to the medial compartment which is typically more affected. These findings demonstrate that cartilage specific loss of OPA1 enhances the severity of age-associated OA in mice.

**Figure 10.**
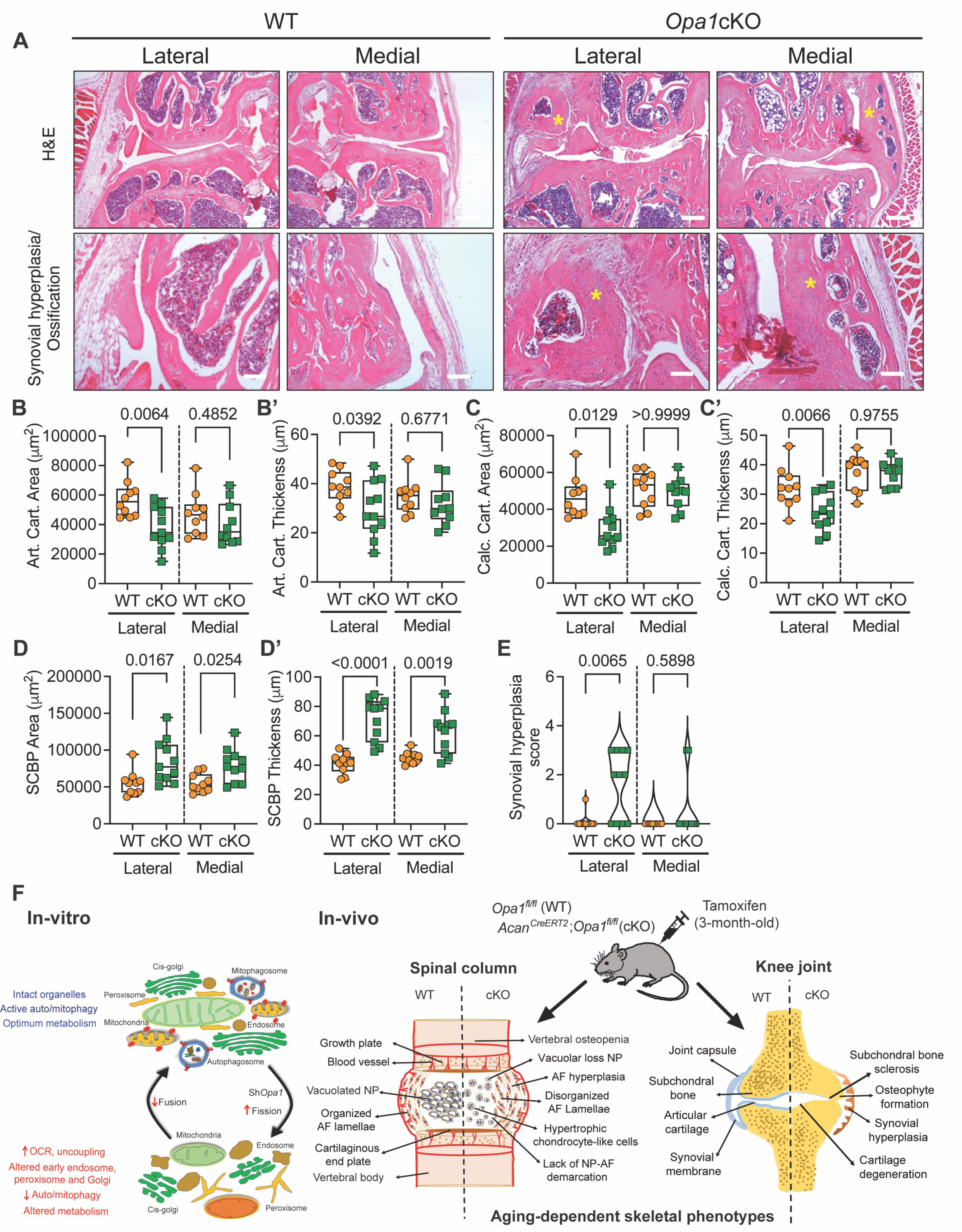
*Opa1*cKO mice evidence synovial hyperplasia and morphological changes in articular cartilage and subchondral bone. (A) Representative H&E images of WT and *Opa1*cKO mouse joint sections showing medial and lateral compartments. Profound synovial hyperplasia and/or ossification is noted in *Opa1*cKO mice in both the medial and lateral compartments (yellow asterisks), when compared to WT mice. Scale bar: Top row - 250 μm, bottom row - 100 μm. (B-E) Histomorphometric analyses of lateral and medial joint compartments conducted on midcoronal sections of WT and *Opa1*cKO mouse limbs. (B, B’) articular cartilage (Art. cart) area and thickness, (C, C’) calcified cartilage (Calc. cart) area and thickness, and (D, D’) subchondral bone plate (SCBP) area and thickness (E) quantitative analysis of synovial hyperplasia. n = 10 WT (4F, 6M), n = 11 *Opa1*cKO (5F, 6M). Quantitative data is represented as box and whiskers (B-D’) or violin (E) plots showing all data points with median and interquartile range and maximum and minimum values. Significance was determined using t-test or Mann Whitney test as appropriate, as appropriate. (F) Schematic showing the in vitro consequences of OPA1-deficiency on mitochondrial and organelle morphology and metabolic functions of NP cells and in vivo phenotypic manifestations of OPA-deletion on the spinal column, and knee joint in *Opa1*cKO mice.

## Discussion

This study for the first time addresses the importance of mitochondria and the IMM protein OPA1 in the maintenance of the health of the spine and knee joint cartilage of aging mice. The reliance on mitochondrial function was surprising since hypoxic chondrocytes and in particular NP cells of IVD primarily rely on glycolysis for energy production (*6, 7*). Relevant to this function, we have previously shown the existence of mitochondrial networks in NP cells (7) and that mitochondrial activity is governed by the HIF-1α-BNIP3 axis controlling mitophagic flux (*6*). We also demonstrated that deletion of mitophagy receptor BNIP3 in NP cells caused mitochondrial dysfunction affecting bioenergetics, triggering meta-inflammation, and resulting in early disc degeneration in mice (*7*). Herein, we show that, OPA1 controls the morphology of mitochondria and cristae as well as multiple organelles including peroxisomes, early endosomes, and cis-Golgi, and that OPA1-loss results in dysregulated autophagy in NP cells. Further, we demonstrate that *Opa1^AcanCreERT2^*mice evidence accelerated age-dependent IVD degeneration, vertebral osteopenia, and severe osteoarthritis of knee joints. Our findings highlight the fact that dysregulation of mitochondrial dynamics affects the metabolism, organelle integrity, and the autophagic/mitophagic pathway causing disc and cartilage degeneration in mice.

OPA1 is required for IMM fusion as well as the maintenance of cristae shape; *Opa1* mutations are linked to multiple human pathologies including vision impairment (*30, 31*), developmental delay, muscle-related disorders, peripheral neuropathy, and cardiomyopathy (*32*). Concerning diseases that afflict the human skeleton, we have previously linked mitochondrial function with spine disease by showing that changes in mitochondrial morphology impact NP cell metabolism and mitophagy (*6, 7*). Herein we observed that deletion of OPA1 resulted in a reduction in mitochondria size but an increase in numbers. This change was unlikely due to mitochondrial biogenesis since there was no difference in the mtDNA content of the mutated cells. (*33–37*). More than likely, as the OPA1-deficient NP cells completely lacked cristae or exhibited detached cristae it indicated that expression of the gene was required for IMM invagination and cristae development. Moreover, since mitochondria are mobile structures, they interact with other organelles and maintain cellular homeostasis while maintaining mitochondrial function (*38*). From this perspective, the observation that the knockdown of OPA1 impacted peroxisomes, endosomes, and cis-Golgi morphology suggested that OPA1 is critical for preserving multi-organelle morphology in NP cells. To the best of our knowledge, this is the first report on the involvement of OPA1 in sustaining the morphology of these organelles.

When mitochondrial quality control was assessed, a significant decrease in key autophagy-related proteins p62 and LC3-II in OPA1-silenced cells was noted. Moreover, bafilomycin A1 treatment showed a lack of LC3-II accumulation suggesting impaired autophagosome formation. LC3-II targets ubiquitin-positive cargoes to sequester them into developing autophagosomes (*39*). Since these autophagy proteins are subjected to regulatory post-translational phosphorylation: mTORC1 (negative) and ULK1 and AMPK (positive).

AMPK promotes autophagy by phosphorylating ULK1 when the ADP/ATP or AMP/ATP ratio is elevated. Indeed, our widely targeted metabolomic measurements indicated that decreased AMP/ATP and ADP/ATP ratios would be associated with decreased phosphorylation, and, as a consequence, reduced autophagy. In contrast to previous studies demonstrating enhanced mitophagy in OPA1 defective cells (*35, 37, 40–42*), we noted that OPA1-knockdown not only inhibited selective autophagy (mitophagy) but also macroautophagy, an event that influenced the morphology of multiple organelles, including the ER, endosomes, and Golgi (*43–45*). For example, enlarged early endosomes and fragmented Golgi, together with autophagosome accumulation are implicated in neurodegenerative diseases such as Down syndrome, Parkinson’s, and amyotrophic lateral sclerosis (*46–48*). Importantly, changes in these organelles underscore the defects in autophagy and disruptions in endocytic and secretory pathways in OPA1-deficient NP cells. Together, our data supports the hypothesis that OPA1 regulates organelle morphology and autophagic/endocytic/secretory pathways in NP cells. One important outcome of these findings is that autophagy and mitochondrial dysfunction are associated with aging, disc degeneration, and the pathogenesis of OA (*3, 49, 50*).

Considering the relationship between OPA1 and mitochondrial function, it is known that in many cell types, which primarily rely on oxidative ATP generation, structural changes in mitochondria profoundly alter energy metabolism. However, since NP cells are predominantly glycolytic, it was important to determine whether alterations in mitochondrial shape and cristae morphology influenced NP cell bioenergetics. We found that while fragmented mitochondria with aberrant cristae morphology consumed more oxygen, it did not manifest in increased oxidative ATP production rates. The number of ATP molecules generated for each dioxygen molecule consumed might vary depending on the mitochondrial efficacy (*51, 52*), suggesting mitochondrial dysfunction in the Sh*Opa1* transduced NP cells. Our findings also differed from prior work showing lower OCR and oxidative ATP generation in OPA1-deficient cells (*36, 53, 54*) but were in agreement with the increase in glycolytic ATP production rate noted in OPA1- deficient neutrophils (*55*). Overall, the results of the current study underscored the observation that OPA1 deficiency influences the energy metabolism of NP cells.

In addition to bioenergetics, the mitochondrion is also the primary site for biosynthetic pathways. Metabolic systems in OPA1-deficient NP cells that were most negatively impacted involved spermidine and spermine biosynthesis and betaine metabolism. Likewise, there was a decrease in amino acid metabolism, taurine and hypotaurine synthesis, pyrimidine, pantothenate and CoA biosynthesis, and urea cycle metabolite. In agreement with these findings, OPA1 deletion in MEF cells resulted in decreased levels of metabolites associated with spermidine and spermine and taurine and hypotaurine pathways (*56*). Notably, spermidine and taurine levels are shown to decline with age, and mitochondrial dysfunction is one of the primary contributors to their deficiency (*57, 58*). Supplementation of spermidine and taurine has been proven to prolong longevity in mice via boosting autophagy/mitophagy, mitochondrial biogenesis, and mitochondrial respiration (*57, 58*). Interestingly, the increased levels of glucose-6-phosphate was, not attributable to enhanced glycolysis associated with the lack of corresponding increase in intracellular and extracellular lactate levels. Depending on cellular metabolism, glucose-6- phosphate shuttles between glycolysis, pentose phosphate shunt, hexosamine biosynthetic pathways, gluconeogenesis, and de-novo lipid synthesis (*59*). Accordingly, in Sh*Opa1* cells, glucose-6-phosphate was utilized for the synthesis of NADP, nucleotide sugar metabolism, and gluconeogenesis pathway activity. Furthermore, elevated malate and the increased malate-aspartate shuttle activity are linked to gluconeogenesis, while increased serine levels in Sh*Opa1* cells indicated de novo serine synthesis, implying that malate was exported to the cytosol (*60, 61*). Interestingly, acetylcarnitine level was decreased in OPA1-deficient cells, which links mitochondrial metabolism to histone acetylation and lipogenesis (*62*). In agreement with a previous report showing OPA1 regulation of lipid metabolism, we also observed reduced fatty acid metabolism, (*63*). OPA1 deficiency also affected one-carbon metabolism with increased serine biosynthesis but decreased utilization in Sh*Opa1* cells. This reduction was likely due to a decrease of s-adenosyl homocysteine and s-adenosyl methionine co-substrates, which are involved in transferring methyl groups for the synthesis of DNA, amino acids, and polyamines. This was further supported by reduced pyrimidine metabolism, polyamines, and amino acids metabolism. Notably, serine biosynthesis and one-carbon metabolism are linked to a variety of mitochondrial disorders (*64*). In summary, the metabolite profiling studies revealed dysregulated metabolism in OPA1-deficient cells.

^13^C labeling experiments utilizing two stable isotope tracers [1,2]-^13^C-glucose and U^13^C- glutamine shed further insights into metabolic dysregulation of OPA1-deficient NP cells. The use of [1,2]-^13^C-glucose MFA revealed a reduction in PDH flux despite an increase in alanine ∑ mn indicating an elevated pyruvate pool. We also noted a decrease in intracellular M+1 glutamate enrichment, an indirect measurement for alpha-ketoglutarate, suggesting pyruvate is prevented from entering the TCA cycle via the conventional pathway. Furthermore, since there is no change in PC flux supports the anaplerosis that pyruvate carboxylation replenishes TCA intermediates and allows the Krebs cycle to continue since the intermediates are not only important for macromolecule synthesis but also for protein post-translational modifications, chromatin modification, and DNA methylation (*65*). Interestingly, in solid hypoxic tumors, increased anaplerosis through PC is required for extracellular collagen production and aberrant fibrosis by tumor-associated fibroblasts, a phenotype we have noted in the discs of *Opa1*cKO mice (*66*). Regarding glutamine utilization, the labeled glutamine entered the TCA via an anaplerotic reaction to alpha-ketoglutarate with M+4 labeling of the forward TCA cycle intermediates succinate, fumarate, malate, and citrate. In contrast to glucose, glutamine labeling showed decreased TCA cycle intermediates such as citrate, succinate, fumarate, malate, and oxaloacetate (aspartate) enrichment. The Citrate M+4 label signifies the oxidative (forward) TCA cycle while the M+5 label is for the reductive (reverse) TCA cycle. We noted a substantial reduction in citrate M+5 label (∼2.5:1 M+4:M+5 citrate), indicating a modest reversal in TCA flux. Of interest, under normal physiological conditions, the majority of succinate is generated by oxidative or forward TCA cycle. Both hypoxia and ETC inhibition are linked to increased levels of succinate. Recent studies found that a part of succinate was generated from fumarate under hypoxia or when ETC was inhibited, implying that fumarate reduction to succinate may act as a valve for surplus electrons from the ETC. Further, it was reported that fumarate was catalyzed explicitly by complex II, rather than passively collecting leaky electrons from the ETC (*26*).

Interestingly, our ^13^C MFA data showed increased succinate ∑ mn compared to fumarate in both ShCtrl and Sh*Opa1* cells, suggesting an oxidative TCA cycle. Furthermore, when we assessed succinate oxidation (fumarate M+4/succinate M+4) and fumarate reduction (succinate M+3/fumarate M+3), we observed no changes in the oxidative process. Overall, these findings suggested that NP cells do not experience metabolic stress by their physiologically hypoxic niche. The *in vitro* studies suggested that mitochondrial morphology and cristae architecture are not only central to energy metabolism, but the integrity of these structures is critical for the proper functioning of mitochondria.

Finally, we investigated the *in vivo* function of OPA1 in the IVD and knee cartilage. We found that *Opa1c*KO mice evidenced enhanced IVD degeneration with aging. Interestingly, the caudal discs showed a more pronounced phenotype than the lumbar discs. Previous studies have shown that caudal discs are more prone to metabolic dysregulation and subsequent degeneration with aging (*67*). Moreover, the caudal spine experiences relatively lower axial loading and different motions than the lumbar spine, it is therefore not unreasonable to hypothesize that the unique interactions between genetics and environmental factors produce different phenotypic outcomes across different spine regions (*68*). Similar to IVD degeneration, aged *Opa1*cKO displayed enhanced age-associated OA severity when compared to control mice; the disease state was characterized by enhanced cartilage damage, osteophyte formation, subchondral bone sclerosis and synovial hyperplasia and/or ossification. usually the medial compartment of the knee is usually most prone to cartilage degeneration (*69*), the lateral knee compartment in *Opa1*cKO mice exhibited significantly more pronounced OA. Similar to caudal discs, the lateral knee compartment experiences lower compressive loads (*70*). Moreover, consistent with previous studies that demonstrate exercise and/or loading in human and animal models can partially restore the functionality of faulty mitochondria (*71, 72*), lack of substantial degeneration in lumbar discs and the medial compartment of the knee suggests a protective effect of loading on these joint tissues (*68, 73*). Of course, in hypoxic skeletal tissues, chronic metabolic stress that reflects abnormalities in mitochondrial function especially over long time period would be expected to influence tissue function especially in aging individuals (*67*).

It is important to note that similar to the NP and knee cartilage, significant degenerative changes affect the AF, underscoring the important contribution of mitochondrial activity to annulus tissue health. This observation is supported by a previous reported microarray analysis of human AF tissues that showed altered genes related to mitochondrial function during degeneration. Moreover, similar to OA development, in degenerated AF tissues, ROS-related gene expression is dramatically altered, suggesting that mitochondrial dysfunction and ROS production promote AF degeneration (*74*). These *in vivo* findings coupled with our mechanistic in vitro studies suggest that the mouse IVD, spinal column, and joint cartilage phenotypes are the results of metabolic dysregulation and cumulative degenerative processes driven by OPA1 deletion. Moreover, in many soft tissues, age-related pathologies, and metabolic disorders due to the accumulation of faulty mitochondria has been exhaustively demonstrated (*75, 76*). From this perspective, targeting and modifying the autophagic pathway and preserving mitochondrial function should be of major concern when designing therapeutics to treat diseases linked to degenerative musculoskeletal conditions.

## METHODS

### Cell isolation, treatments, and hypoxic culture

Primary NP cells from Sprague Dawley (Charles River), were obtained and cultured in antibiotic supplemented Dulbecco’s modified Eagle medium (DMEM) and 10% FBS (*77*). To explore role of OPA1, lentiviral particle production and viral transduction LV-Sh*Opa1* clone #1 (TRCN0000091111) and LV-SH*Opa1* clone #2 (TRCN0000348537) (Sigma, St. Louis, MO, USA) and pLKO.1 ShCtrl and psPAX2 (#12260) and pMD2G (#12259) (Addgene, Cambridge, MA, USA) were used. 1 day before transfection, HEK 293T cells were plated in 10-cm plates (5x10^6^ cells/plate) in DMEM with 10% heat-inactivated FBS. ShCtrl or Sh*Opa1* plasmids, as well as psPAX2 and pMD2.G, were transfected into cells and lentiviral particles were collected 48 to 60 hours after transfection. Cells were transduced with viral particles using 8 mg/ mL polybrene and after 3 days of transduction cell were cultured hypoxia workstation (Invivo2 400; Baker Ruskinn, Bridgend, UK) with a mixture of 1% O2, 5% CO2, and 94% N2 for 24 h and day 4 cells were harvested for protein extraction. Because both OPA1 shRNA clones had equal effects on mitochondrial morphology and mitophagy, cells transduced with LV-Sh*Opa1* #1 (TRCN0000091111) were utilized for all metabolic investigations.

### Immunocytochemistry

ShCtrl, Sh*Opa1* or DFP treated NP cells were plated on glass coverslips and treated with 100 nM MitoTracker Red CMXRos (Thermo Fisher Scientific, Waltham, MA, USA; M7512) for 30 minutes after completion of the experimental treatments. Cells were then fixed with 4% PFA or ice-cold methanol for 15 minutes and permeabilized with 0.1% Triton X-100 for 10 minutes and blocked with 1% BSA for 1 h. Cells were incubated with anti-OPA1 (612607) (BD Bioscience), anti-PMP70 (PA1-650), anti-GM130 (MA5-35107) (Fisher Scientific, Pittsburgh, PA, USA), anti-EEA1 (24115), anti-BNIP3 (3769S), anti-LC3B (12741) (Cell signaling, Denver, MA, USA), anti-TNG46 (ab16059), anti-LAMP1 (ab24170), and anti-BNIP3L (ab109414), (Abcam, Cambridge, MA, USA), and anti-RAB7 (R8779) (EMD Millipore, Burlington, MA, USA), in blocking buffer at 1:100 to 200 at 4 °C overnight. After washing, cells were incubated with Alexa Fluor 488 and mounted with ProLong Gold Antifade Mountant with DAPI. To confirm the specificity of staining cells were incubated with isotype mouse (7076P2) or rabbit (7074P2) IgG antibodies (Cell Signaling, Danvers, MA, USA) (Fig. S7). Cells were visualized using a Zeiss LSM 800 Axio Inverted confocal microscope (Plan-Apochromat 40x/1.3 oil or 63x/1.40 oil) for positive staining for markers OPA1, LC3, PMP70, EEA1, GM130, TGN46, RAB7, LAMP1, BNIP3 and BNIP3L.

### Organelles morphology analysis

Mitochondria, peroxisomes, endosomes and Golgi number, branching and morphology were quantified in ImageJ using methods reported earlier (*6, 7*). Briefly, the confocal images were converted to binary by threshold and then converted to a skeleton that represented the features in the original image using a wireframe of lines one pixel wide. All pixels within a skeleton were then measured using analyze skeleton. The output will give the number of particles which denotes the total number of mitochondria, the aspect ratio (AR) represents the “length to width ratio” and the form factor (FF), the complexity and branching aspect of mitochondria were calculated from circularity. For the endosome, Feret diameter was used to plot the graph.

### Western Blotting

ShCtrl, ShOpa1 transduced cells were lysed and 35 μg of protein was electroblotted to PVDF membranes (EMD Millipore, Burlington, MA, USA). The membranes were blocked and incubated overnight at 4°C with antibodies against anti-BNIP3 (3769), anti-BNIP3L (12396) , anti-LC3B (2775) , anti-LAMP1 (9091) , anti-p62 (5114), anti-CHOP (78063), anit-Beclin1 (3738), anti-MFF (84580), anti-MFN2 (9482), anti-FIS1 (32525), anti-Ub (#), pUb (62802), anti-GAPDH (5174) (Cell Signaling), anti-DRP1 (611113), anti-OPA1 (612607) (BD Biosciences, San Jose, CA, USA), anti-PARKIN (sc-32282), anti-MFN1 (ab126575) (Abcam, Cambridge, MA, USA), Immunolabeling was detected using an ECL reagent (Azure biosystems 300, Dublin, CA, USA). The membranes were detected using ECL reagent (Azure biosystems, Dublin, CA, USA). Densitometric analysis was performed using ImageJ software.

### Seahorse XF analysis

Maximum glycolytic capacity and ATP production rate using methods reported by Mookerjee and colleagues (*7, 24, 25*). In brief, ShCtrl, Sh*Opa1* cells were plated in a 24-well Seahorse V7- PS test plate under hypoxia 24 hours before the experiment. Cells were washed three times with 500 μl of KRPH (Krebs Ringer Phosphate HEPES) before being cultured for one hour at 37 °C in 100% air. For glycolytic capacity calculation, oxygen consumption rate (OCR) and related extracellular acidification rate (ECAR) were determined in a Seahorse XFe24 analyzer (Agilent Techonoligies) by adding 10 mM glucose, 1 μM rotenone plus 1 μM myxothiazol, and 200 μM monensin plus 1 μM FCCP via ports A-C. OCR and ECAR were assessed by adding 10 mM glucose, 2 μg oligomycin, 1 μM rotenone plus 1 μM myxothiazol to determine ATP generation rate from oxidative and glycolytic pathways.

### Widely targeted small metabolites measurements

Cells were transduced with ShCtrl and ShOpa1 viral particles as described above. On the third day, the medium was changed to DMEM without pyruvate, 10% dialyzed FBS (Sigma F0392), and the cells were grown under hypoxia for 24 hours. Cells were washed and collected in ice-cold 80% methanol before being snap-frozen in liquid nitrogen and kept at -80 degrees Celsius until use. Prior to analyzing the metabolites, cell pellet samples were centrifuged and pipetted into a LC sampling vial. Each sample had internal standards. After drying under mild nitrogen flow, the samples were reconstituted in 150 μl of 80% methanol for injection. The samples were analyzed on an ABsciex 6500+ coupled with a Waters UPLC. Small metabolites were separated using the Ace PFP column and the iHILIC-p column (HILICON) and a pooled quality control (QC) sample was added to the sample list. The QCs sample was injected six times to calculate the coefficient of variation (CV) for data quality control. Metabolites with CVs lower than 30% used for the quantification. MetaboAnalyst 5.0 web server was used to analyze the data, and acceptable metabolites were manually input using the HMDB number. The small metabolites pathway data bank (SMPDB), which contains 99 compounds based on normal human metabolic pathways, was used for enrichment and pathway analysis. MetaboAnalyst provides the list of pathways in which these metabolites are found.

### 13C-Metabolic flux analysis

For [1,2]-^13^C-glucose flux analysis, 50% of DMEM media contained ^13^C-labeled glucose. A volume of 100 μl of cell culture medium from 1,2-^13^C glucose experiment was treated with 400 μl of methanol. After centrifugation, the supernatant was transferred to a LC-MS sampling vial and dried under gentle nitrogen flow. The sample was reconstituted into 100 ml of 80% methanol for LC-MS injection. Metabolite separation was performed on an ACE PFP-C18 column (1.7 μM x 1 mm x 100 mm) and analyzed on a ABSciex 6500+ with a Multiple Reaction Monitoring (MRM) mode. The glycolysis, pentose cycle, PDH, PC, PDH/PC, and PDH+PC fluxes were calculated using methods reported by Madhu et al., (*7*).

When flux was assessed in the U^13^C-glutamine labeling experiment, labeled glutamine was added to be 50% of the total DMEM glutamine concentration. Methanol extraction from [1,2]-^13^C-glucose and U^13^C glutamine labeled cell pellets were dried under gentle nitrogen flow. The dried samples were derivatized with a methyl-moximation (with 15 mg/ml methoxy amine in pyridine, 30 °C for 90 minutes) and MTBSTFA (at 70°C for 60 minutes). The samples were then analyzed with an Agilent GC-MS, with an electron impact mode and a DB-5MS column (Agilent) following our protocol (*7*). The data were analyzed with Mass Hunter Quantitative Analysis software (Agilent). The enrichment was calculated after subtraction from the background of the non-labeled treatment samples. Fractional enrichments from G to R is sigma (stands for sigma mean), and it is equal to the weighted mean average of the metabolite’s enrichment (sigma mn = 1 xm1+2xm2+3xm3, etc).

### Mitochondrial DNA quantification

DNA was isolated from ShCtrl and Sh*Opa1* cells using DNA extraction lysis buffer (Qiagen). The mitochondrial DNA (mtDNA) content was determined using qPCR (Applied Biosystems A25742 PowerUP SYBR green master mix). For mitochondrial DNA (mtDNA; Nd1) 5’-GGC TCC TTC TCC CTA CAA ATA C-3’ and 5’-TGT TTC TGC AAG GGT TGA AAT G-3’. For nuclear DNA (nDNA; Cox4) 5’-ATG TTG ATC GGC GTG ACT AC-3’ and 5’-AGT GGGCCT TCT CCT TCT-3’ were used. The ratio of mtDNA to nDNA (mtDNA/nDNA) reflects the relative mtDNA content.

### Generation of conditional knockout mice

All mouse studies were carried out in conformity with the Institutional Animal Care and Use Committee (IACUC) of Thomas Jefferson University’s applicable rules and regulations. *Opa1^fl/fl^* on mixed C57BL/6–129/SvEv background were described previously (*54*). Aggrecan-CreERT2 mice were from Jackson Laboratories (stock #019148). OPA1 conditional knock-out (*Opa1*cKO:*Acan^CreERT2^Opa1^fl/fl^*) and control (*Opa1*CTR: *Opa1^fl/fl^*) mice were generated and analyzed after 3 (7-month-old), 9 (12-month-old), and 17 (20-month-old) months after deletion to determine degree of degeneration. In brief, *Opa1^fl/fl^* mice have loxP sites that flanking exon 10-13 of the *Opa1* gene, which encodes essential GTPase domain. Cre-mediated recombination of these alleles inactivates OPA1 in target cells. In addition, upon loxP recombination, a stop codon was generated immediately after exon 9 due to a frame shift. For all experiments, skeletally matured 3-month-old female and male mice of all genotypes received an intraperitoneal injection of 100 mg/kg tamoxifen (Sigma-Aldrich, St. Louis, MO, USA) dissolved in palm oil (Sigma-Aldrich) for 3 consecutive days to activate Cre recombinase.

### Histological analysis of intervertebral disc

Spines were dissected and immediately fixed in freshly made 4% paraformaldehyde (PFA) for 48 hours, followed by decalcification in 20% EDTA at 4°C prior to embedding in paraffin.

Lumbar and caudal motion segments at 7 months (n=4-5 lumbar discs/animal, 2-3 caudal discs/animal, 6-7 animals/genotype, 25-30 lumbar and 12-14 caudal discs/genotype) and 12 months (n=4-5 lumbar discs/animal, 2-3 caudal discs/animal, 8-9 animals/genotype, 32-36 lumbar and 18-19 caudal discs/genotype) were processed in addition to lumbar and caudal motions segments also 7, 12 and 20 months (n=4 lumbar discs/animal, 3 caudal discs/animal, 9-11 animals/genotype, 36-44 lumbar and 27-33 caudal discs/genotype). Coronal sections 7 μm in thickness were stained with 1% SafraninO, 0.05% Fast Green, and 1% Hematoxylin to assess morphology and imaged on an Axio Imager A2 microscope using 5x/0.15 N-Achroplan or 20x/0.5 EC Plan-Neofluar (Carl Zeiss) objectives, Axiocam 105 color camera, and Zen2™ software (Carl Zeiss AG, Germany). To evaluate disc degeneration, 4 blinded observers used a Modified Thompson grading scale for the NP and AF perform histological scoring of 7, 12- and 20-month-old WT and Opa1cKO animals. Higher scores reflect a higher degree of degeneration. Picrosirius red staining (Polysciences, 24901) was used to measure collagen fibril thickness, and pictures were captured using a polarizing light microscope (Eclipse LV100 POL; Nikon) and a 4x /0.25 Pol /WD 7.0 (Nikon) objective. The color thresholds for green (thin), yellow (middle), and red (thick) fibers were established using the Nikon NIS Elements Viewer program (*78, 79*). For all samples, the color threshold values remained constant.

### Micro-computed tomography (μCT) analysis of mouse lumbar spine and hindlimb

μCT imaging was performed on the lumbar spine of 7, 12, and 20-month WT and Opa1cKO mice (n = 4 lumbar discs/animal, 9 animals/genotype, 52-54 lumbar discs/genotype) using the high-resolution μCT scanner (Skyscan 1272, Bruker, Belgium). Lumber segments L2-6 were placed in PBS and scanned with an energy of 50 kVp and current of 200 mA resulting in 15 mm^3^ voxel size resolution. Trabecular parameters were assessed in the 3D reconstructed trabecular tissue using Skyscan CT analysis (CTAn) software by contouring the region of interest (ROI).

The bone volume percentage (BV/TV), trabecular number (Tb. N.), trabecular thickness (Tb. Th.), and trabecular separation (Tb. Sp.) of the resulting datasets were all evaluated. Cortical bone volume (BV), cross-sectional thickness (Cs. Th.), mean cross-sectional bone area (B. Ar), and mean cross-sectional tissue area (T. Ar) were all measured in two dimensions. A standard curve was established with a mineral density calibration phantom pair (0.25 g/cm3 CaHA and 0.75 g/cm3 CaHA) to determine mineral density. Intervertebral disc height and the length of the vertebral bones were measured and averaged along the dorsal, midline, and ventral regions in the sagittal plane. Disc height index (DHI) was calculated as previously described (*78, 79*).

Mouse hindlimbs (right limb) were harvested, surrounding muscles were removed, and limbs were fixed in formalin (10%, 48 hours). MicroCT analysis of the femur, tibia, and knee joint were performed on a Bruker SkyScan 1275 scanner as we have previously described (*29*). MicroCT reconstructions and quantitative analysis of tibial subchondral bone volume fraction (BV/TV), trabecular thickness (Tb.Th), trabecular separation (Tb.Sp), and subchondral bone plate thickness (SCBP) were performed using the SkyScan CT Analyzer (CTan) and CT Vox software on coronal slices that spanned the medial and lateral tibial plateaus.

### Histological and histomorphometry analysis of OA in mouse hindlimbs

Hindlimbs were decalcified (EDTA, 19%, 21 days), processed, paraffin embedded, and sectioned along the coronal plane, as we have described (*29*). Mid-coronal sections were stained with hematoxylin and eosin (H&E), or toluidine blue and OA severity was analyzed by Articular Cartilage Structure (ACS), toluidine blue, and osteophyte scoring on the medial and lateral tibial plateaus (MTP, LTP) and femoral condyles (MFC, LFC). Synovial hyperplasia was assessed using a 0-3 scale as described earlier (*80*) In joints that presented with synovial ossification, a maximal synovial hyperplasia score was assigned. Analysis was performed by a blinded scorer with experience of the OA scoring techniques. Detailed histomorphometric analysis of articular cartilage thickness and area, calcified cartilage thickness and area, and subchondral bone thickness and area were analyzed on the MTP and LTP using ImageJ software as we have previously described (*29*).

### Immunohistochemistry and confocal analysis

For immunofluorescence microscopy mid-coronal disc tissue sections of 7 μm thickness were de-paraffinized and incubated in microwaved citrate buffer for 20 min, proteinase K for 10 min at room temperature, or Chondroitinase ABC for 30 min at 37 °C for antigen retrieval (n = 5-6 lumbar discs/animal, 3 animals/genotype, 17-18 lumbar discs/genotype). Sections were blocked in 5% normal serum (Thermo Fisher Scientific, 10000 C) in PBS-T (0.4% Triton X-100 in PBS) and incubated with antibody against GLUT-1 (1:200, Abcam, ab40084), CA3 (1:150; Santa Cruz Biotechnology, sc-50715) Collagen X (1:500, Abcam, ab58632) Collagen I (1:100; Abcam, ab34710), COMP (1:200, Abcam, ab231977), ACAN (1:50; Millipore Sigma, AB1031), ARGXX. (1:200; Abcam, ab3773). For mouse primary antibody Mouse on Mouse Kit (Vector laboratories, BMK2202) was used for blocking and primary antibody incubation. Tissue sections were thoroughly washed and incubated with Alexa Fluor−488 or 594 or 647 conjugated secondary antibodies (Jackson ImmunoResearch Lab, Inc.), at a dilution of 1:700 for 1 h at room temperature in dark. The sections were washed with PBS-T (0.4% Triton X-100 in PBS) and mounted with ProLong® Gold Antifade Mountant with DAPI (Thermo Fisher Scientific, P36934). All mounted slides were visualized using a Zeiss LSM 800 Axio Inverted confocal microscope (Plan-Apochromat 5x/0.16 or 10x/0.45). ImageJ 1.52i (NIH) was used for all quantifications. Images were threshold to generate binary images, then NP and AF compartments were contoured manually using the Freehand Tool. The ROI were examined using the Area Fraction measurement.

### Statistical analysis

Statistical analysis was performed using Prism9 (GraphPad, La Jolla, CA, USA). The quantitative data are represented as mean ± SEM or Box and whisker plots showing all data points with median and interquartile range and maximum and minimum values. Data distribution was checked with Shapiro-Wilk normality test, and differences between two groups were assessed by t-test or Mann-Whitney test as appropriate. One-way ANOVA, whereas non-normally distributed data were analyzed using or Kruskal Wallis test with appropriate post-hoc test (Sidak’s multiple comparisons test) was used for comparisons between more than two groups. Analysis of Modified Thompson Grading data distribution and fiber and fiber thickness distribution were performed using chi-square test; p <0.05.

## ACKNOWLEDGEMENTS

We thank Drs. Irwin Kurland and Yunping Qiu, Stable Isotope and Metabolomics Core Facility, Diabetes Research and Training Center (DRTC) at Albert Einstein College of Medicine for metabolic flux analyses. We thank Dr. Gyorgy Csordas and Timothy Schneider at the Center for Mitochondrial Research Imaging and Diagnostic Facility of Thomas Jefferson University for help with TEM studies.

## AUTHORS’ CONTRIBUTIONS

Conceptualization: VM, MVR; Methodology: VM, MVR; Investigation: VM, MHM, AC, KS, KI, OH, PKB, JC; Visualization: VM; Supervision: MVR; Writing - original draft: VM, MVR; Writing - review & editing: VM, KS, MVR, JC, HS.

## DATA AND MATERIALS AVAILABILITY

All data generated or analyzed during this study are included in this published article and its Supplementary files.

## ETHICS STATEMENT

All animal experiments were performed under IACUC protocols approved by Thomas Jefferson University

## SUPPLEMENTAL FIGURES

Figure S1. **OPA1 knockdown does not alter OMM fusion, fission, and mitophagy related proteins in NP cells.** (A, A’) Western blot analysis and corresponding densitometric quantification of fusion proteins OPA1, MFN1, MFN2 and fission proteins DRP1, MFF, FIS1 in NP cells lentivirally transduced with ShCtrl, and Sh*Opa1* #1 and Sh*Opa1*#2. (B) Immunofluorescence staining for BNIP3 and NIX in ShCtrl and Sh*Opa1*#1 NP cells. Scale bar = 15 μm. (C, C’) Western blot analysis and corresponding densitometric quantification of autophagy markers BECLIN1, ATG12-ATG5, mitophagy markers BNIP3, NIX, and apoptosis marker CHOP. (D, D’) Western blot analysis and densitometric quantification of Ubiquitin in NP cells lentivirally transduced with ShCtrl, Sh*Opa1*#1, and Sh*Opa1*#2. Data represents 6 independent experiments and presented as Box and Whiskers plots showing all data points with median and interquartile range and maximum and minimum values. Statistical significance was determined using one-way ANOVA with Sidak’s multiple comparisons test.

Figure S2. **Effect of OPA-1 deficiency in NP cells on cis-Golgi, trans-Golgi, late endosome and lysosomal morphology.** (A) A representative TEM image of ShCtrl and Sh*Opa1* cells showing fragmented cis-Golgi compared in Sh*Opa1* cells. Scale bar: 200 nm (B) Immunofluorescence staining of ShCtrl and Sh*Opa1* NP cells for Trans-Golgi marker TGN46, late endosome marker RAB7 and lysosome marker LAMP1. There are no distinctive morphological changes in either of these organelles between the two groups. Scale bar: 15 μm

Figure S3. **Metabolic pathway analysis (MetPA)** of (A) upregulated and (B) downregulated metabolites (FDR < 0.05) from Sh*Opa1 vs* ShCtrl transduced cells.

Figure S4. **IVD degeneration phenotype in *Opa1*cKO mice is age-dependent** (A-B) Representative images of Safranin O/Fast Green/hematoxylin staining of (A) 7-month-old and (B) 12-month-old WT and *Opa1*cKO lumbar discs showing disc morphology and overall proteoglycan content in the intervertebral disc, scale bar = 500 μm. (A’ A’, B’ B”) Modified Thompson Scoring (A’, B’) grade distributions and (A”, B”) average scores from NP and AF compartments of 7-month-old and 12-month-old WT and *Opa1*cKO lumbar discs. n = 7-month-old: 6-7 mice/genotype; 4/5 lumbar disc/animal, 25-34 discs/genotype; 12-month-old: 9-10 mice/genotype; 4 lumbar disc/animal, 36-40 discs/genotype. (C) Representative Safranin O/Fast Green/Hematoxylin staining images of 7-month-old WT and *Opa1*cKO caudal discs, scale bar = 500 μm. (C’, C”) Modified Thompson Scoring grade distributions and average scores of NP and AF compartments of 7-month-old WT and *Opa1*cKO caudal discs. (D) Representative Safranin O/Fast Green staining images of 12-month-old WT and *Opa1*cKO caudal discs; scale bar = top row – 500 μm, bottom rows - 100 μm. (D’, D”) Modified Thompson Scoring grade distributions and average scores of NP and AF compartments of 12-month-old WT and *Opa1*cKO caudal discs. (D”’) Quantification of AF hyperplasia in WT and *Opa1*cKO caudal discs. N = 7-month-old: 6-7 mice/genotype; 2 caudal disc/animal, 12-14 discs/genotype; 12-month-old: 9-10 mice/genotype; 2/3 caudal disc/animal, 19-24 discs/genotype. Significance was determined using a chi-square test (A’, B; C’, D’) and data represented as Violin plots or Box and Whisker plot showing all data points with median and interquartile range and maximum and minimum values. Statistical significance was determined using t-test or Mann Whitney test, as appropriate.

Figure S5. *Opa1*cKO mice show mild degenerative changes in lumbar discs at 20-months. (A) Representative Safranin O/Fast Green/Hematoxylin staining images of 20-month-old WT and *Opa1*cKO lumbar discs, scale bar = 500 μm. (A’, A”) Modified Thompson Scoring (A’) grade distributions and (A”) average scores of NP and AF compartments of 20-month-old WT and *Opa1*cKO lumbar discs. (B) Representative picrosirius red stained polarized light images and (B’) quantification of 20-month-old lumbar discs showing the presence of collagen fibers in the NP compartment. n = 9-11 mice/genotype; 4 lumbar disc/animal, 36-44 discs/genotype. Significance for A” and B’ was determined using a chi-square test. Significance for A” was determined using an unpaired t-test or Mann-Whitney test, as appropriate. Quantitative data in A” is represented as a Violin plot showing all data points with median and interquartile range and maximum and minimum values.

Figure S6. ***Opa1cKO mice* did not show altered cartilage morphology at 12-months.** WT and *Opa1*cKO mice were aged to 12 months and OA severity was analyzed by microCT and histological analysis. (A) The representative 3D reconstruction of the knee joint of 12-month-old WT and *Opa1*cKO mice. Scale bar: 500 μm. (B-E) Quantitative microCT analyses of tibial subchondral bone parameters on the lateral and medial tibial plateaus of 12-month-old WT and *Opa1*cKO mice (B) subchondral bone volume fraction (BV/TV), (C) trabecular thickness (Tb.Th), (D) trabecular separation (Tb.Sp), (E) subchondral bone plate thickness (SCBP.Th). (F-G) Representative images of H&E and toluidine blue stained midcoronal sections showing the lateral and medial tibial plateaus and femoral condyles from WT and *Opa1*cKO mice. (H-J) Summed (MTP, MFC, LTP, LFC) scores for (H) Articular Cartilage structure (ACS), (I) toluidine blue and (J) osteophyte in 12-month-old mice. LFC = lateral femoral condyle, LTP = lateral tibial plateau, MFC = medial femoral condyle, MTP = medial tibial plateau. (K, K’) Histomorphometric analyses of lateral and medial compartments performed on midcoronal sections of 12-month-old WT and *Opa1*cKO mouse limbs to compute (K) articular cartilage (Art. cart) area, (K’) articular cartilage (Art. cart) thickness, (L) calcified cartilage (Calc. cart) area and (L’) calcified cartilage (Calc. cart) thickness, (M) subchondral bone plate (SCBP) area and (M’) subchondral bone plate (SCBP) thickness. n = 7 WT (4F, 3M), 9 *Opa1*cKO (4F, 5M). Quantitative data represented as Box and Whiskers plots showing all data points with median and interquartile range and maximum and minimum values. Significance was determined using an unpaired t-test or Mann-Whitney test, as appropriate.

Figure S7. **Specificity of negative controls employed for immunofluorescence staining.** Cells were stained with MitoTracker Red CMXRos and fixed followed by incubation with mouse isotype antibody with anti-mouse secondary Alexa Flour 488 antibody or rabbit isotype antibody with anti-rabbit secondary Alexa Flour 488 antibody and mounted with ProLong Gold Antifade Mountant with DAPI. Isotype antibody-negative controls were used to validate the specificity of immunostaining performed on NP cells.

