## Supplementary figures and images for "OPA1 protects intervertebral disc and knee joint health in aged mice by maintaining the structure and metabolic functions of mitochondria"

### Supplementary figure 1-7

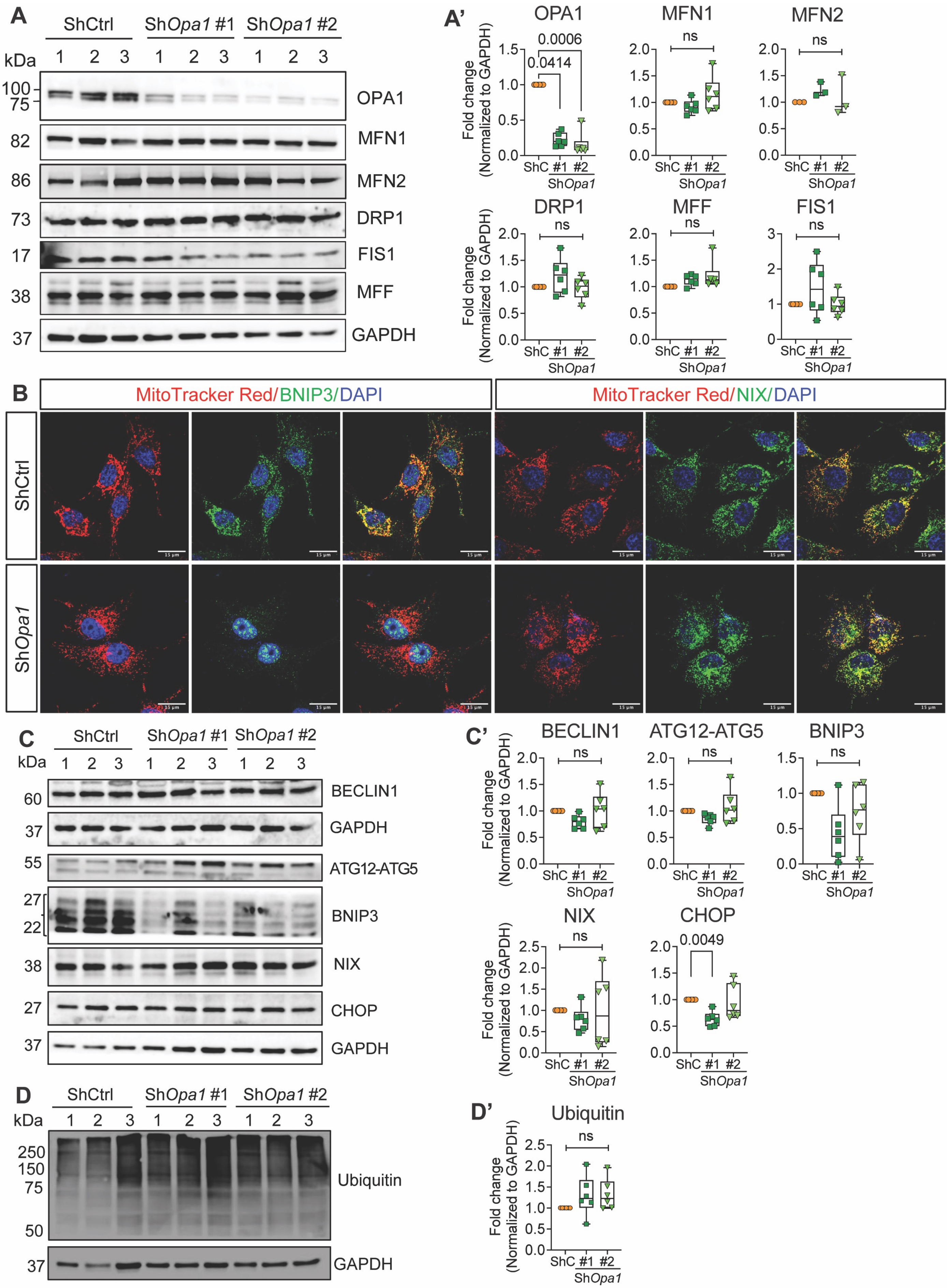

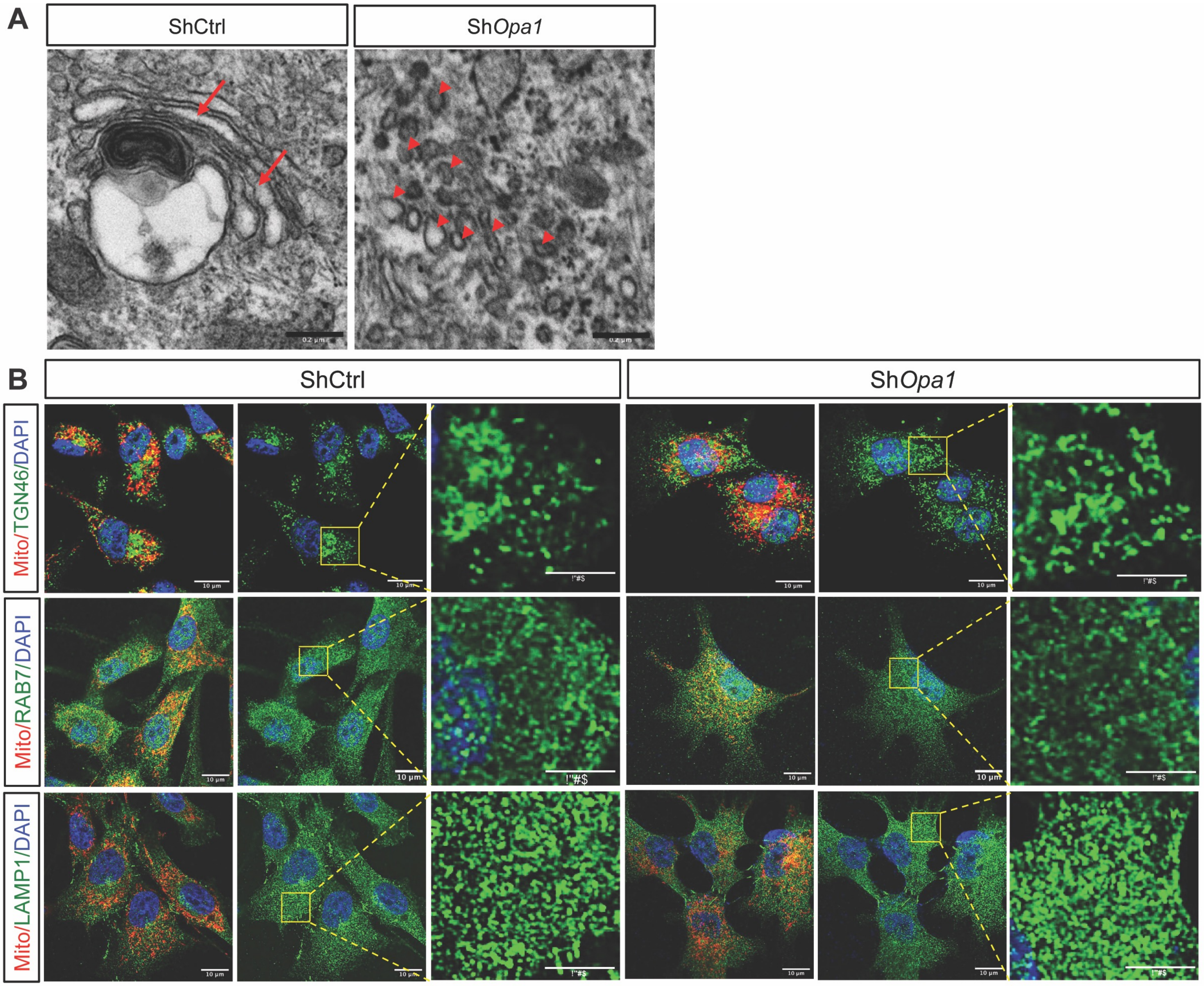

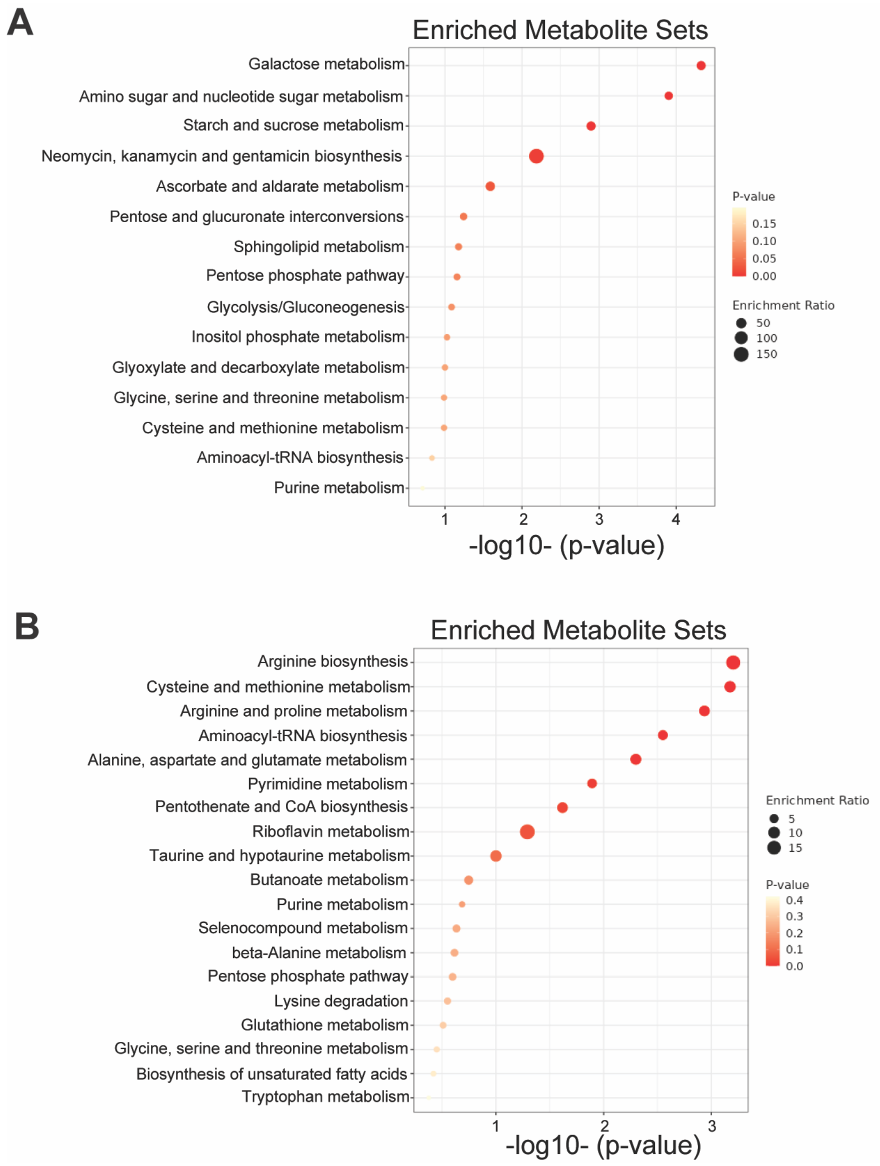

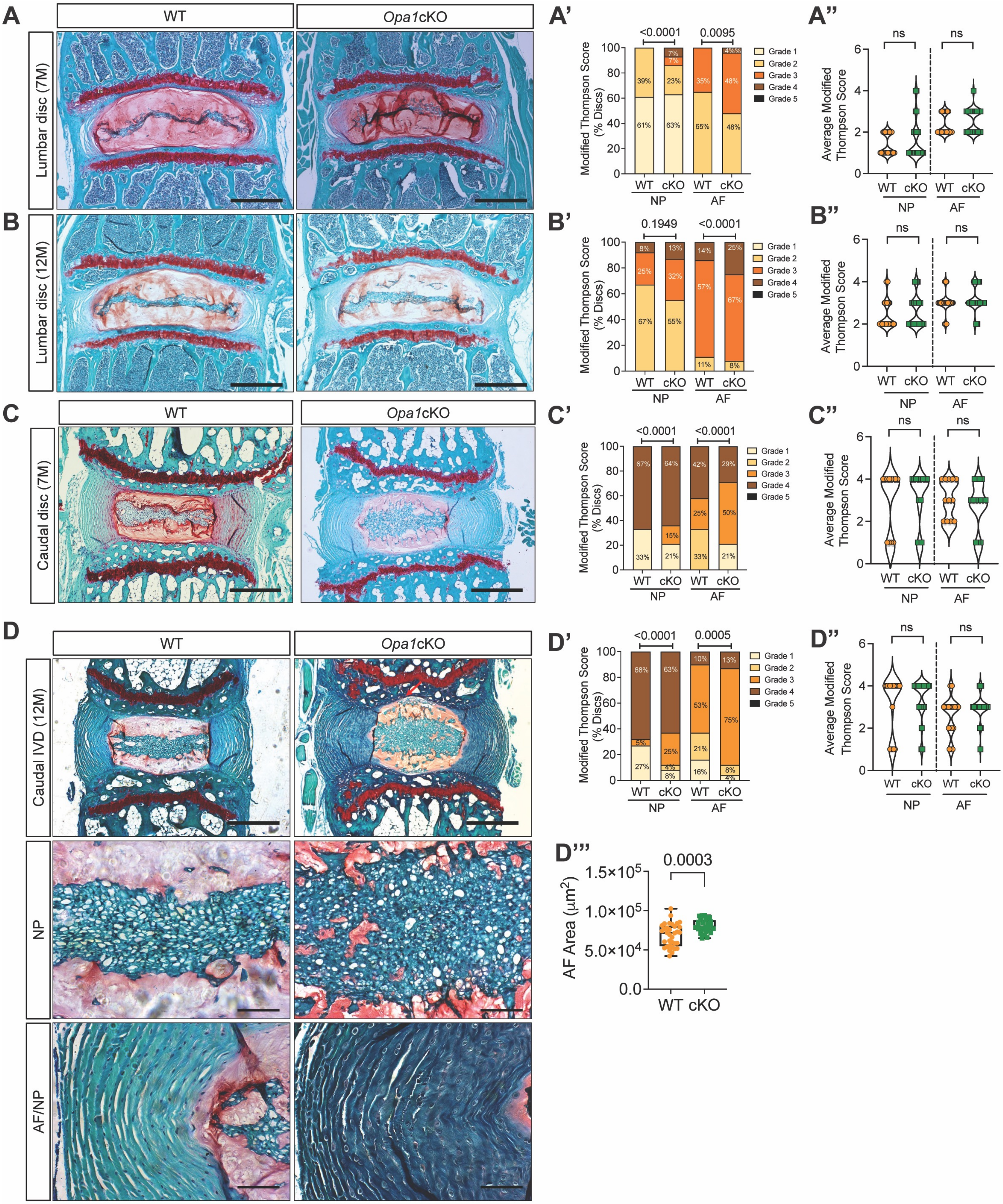

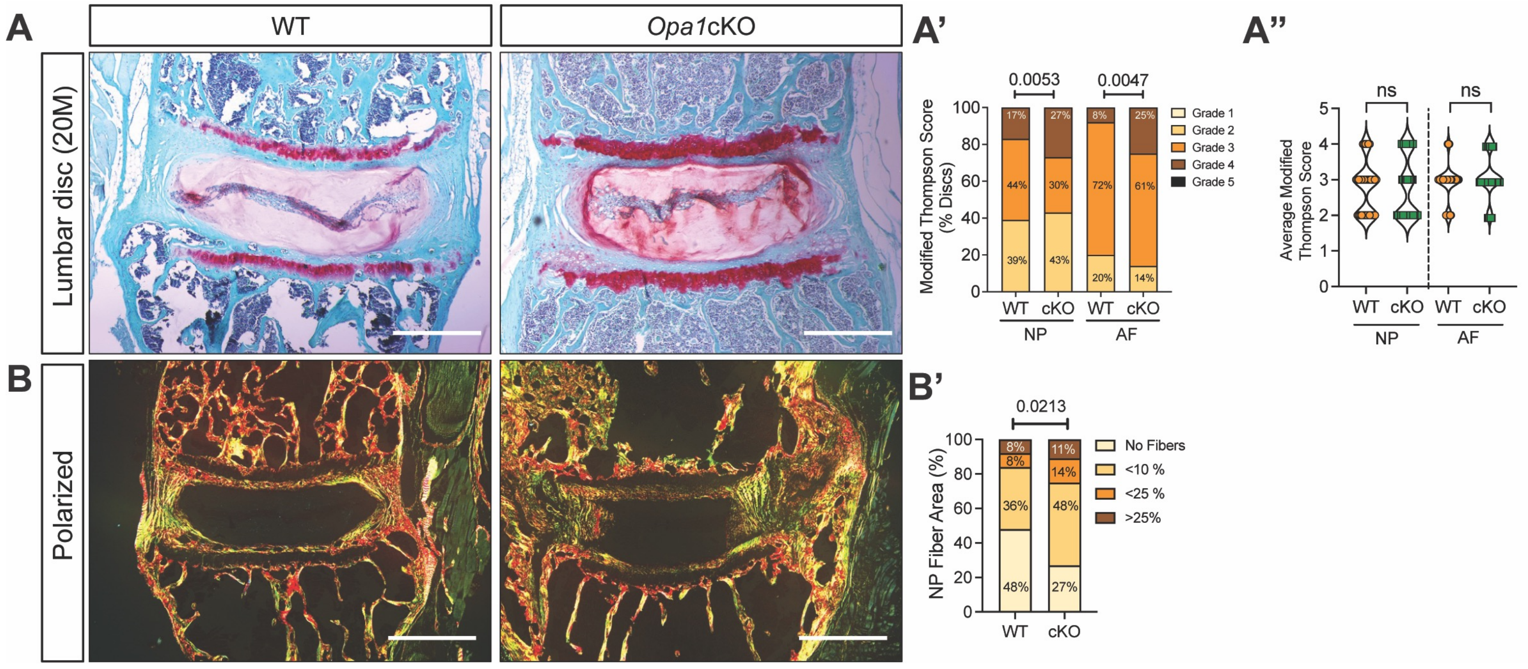

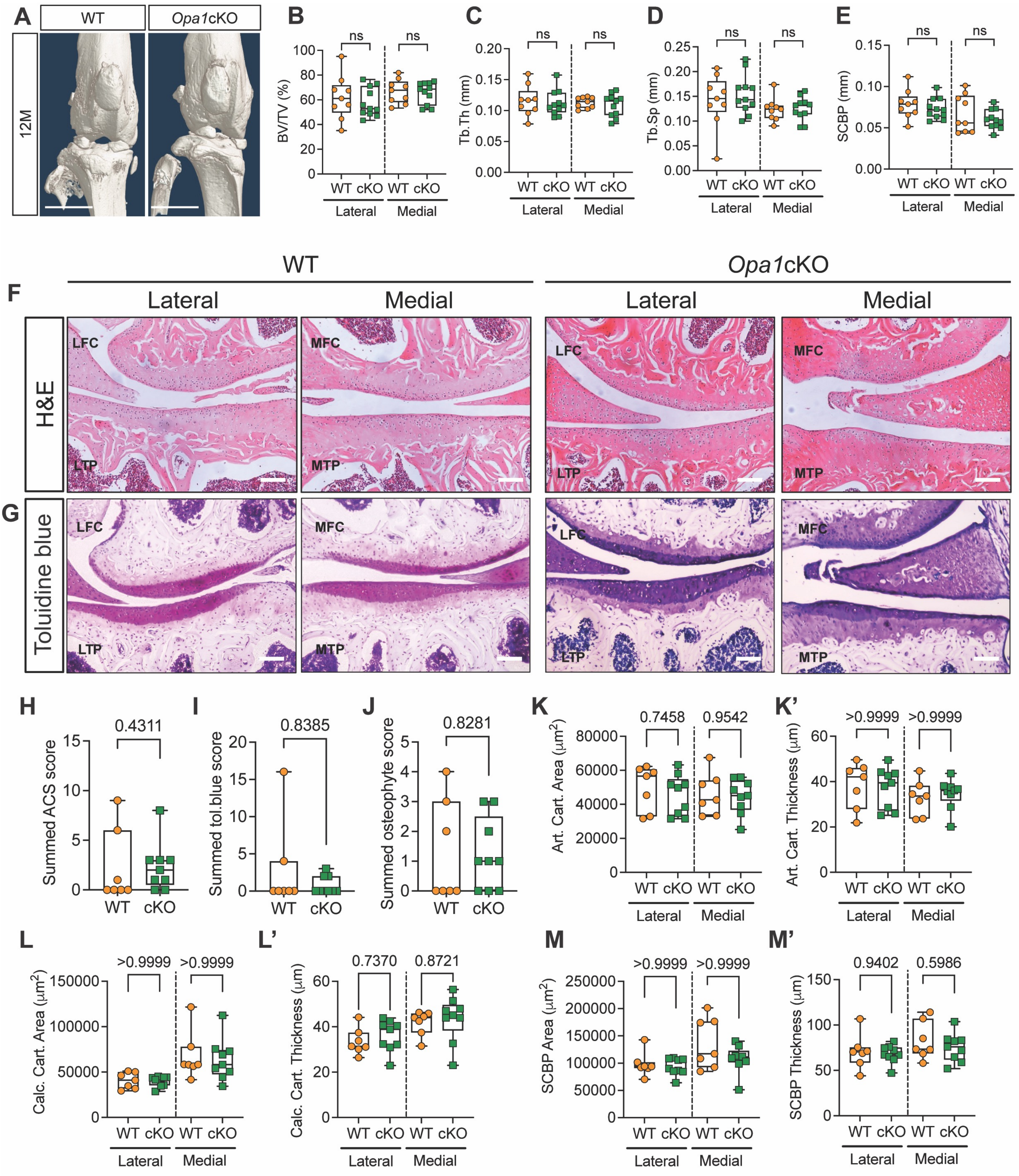

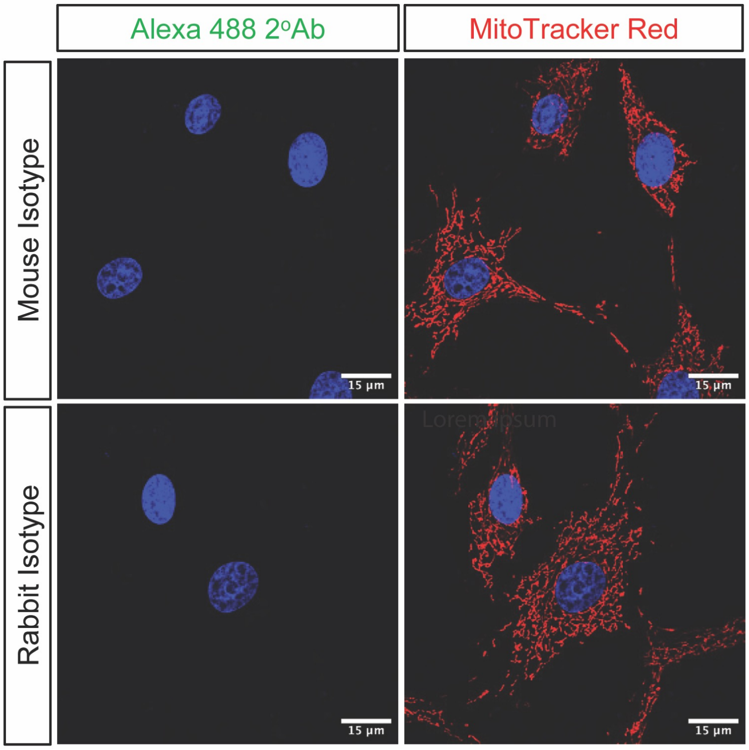
